# Multi-omics reveals principles of gene regulation and pervasive non-productive transcription in the human cytomegalovirus genome

**DOI:** 10.1101/2022.01.07.472583

**Authors:** Christopher Sebastian Jürges, Manivel Lodha, Vu Thuy Khanh Le-Trilling, Pranjali Bhandare, Elmar Wolf, Albert Zimmermann, Mirko Trilling, Bhupesh Prusty, Lars Dölken, Florian Erhard

## Abstract

For decades, human cytomegalovirus (HCMV) was thought to express ≈200 viral proteins during lytic infection. In recent years, systems biology approaches uncovered hundreds of additional viral gene products and suggested thousands of viral sites of transcription initiation. Despite all available data, the molecular mechanisms of HCMV gene regulation remain poorly understood. Here, we provide a unifying model of productive HCMV gene expression employing transcription start site profiling combined with metabolic RNA labeling as well as integrative computational analysis of previously published big data. This approach defined the expression of >2,600 high confidence viral transcripts and explained the complex kinetics of viral protein expression by cumulative effects of translation of incoming virion-associated RNA, multiple transcription start sites with distinct kinetics per viral open reading frame, and differences in viral protein stability. Most importantly, we identify pervasive transcription of transient RNAs as a common feature of this large DNA virus with its human host.

## Main

HCMV is an ubiquitous member of the herpesvirus family and responsible for life-threatening disease in immune-immature, immune-compromised, and immune-senescent individuals ^1–3^. The application of molecular high-throughput techniques greatly expanded the known repertoire of HCMV-encoded RNAs and proteins. In a landmark study using ribosome profiling (Ribo-seq), hundreds of novel viral gene products were identified ^4^. Improved computational analysis refined the viral translatome and identified hundreds of additional viral open reading frames (ORFs) raising the number of HCMV ORFs translated during lytic infection of fibroblasts to >1,000 ^5^.

HCMV genes are regulated in cascades of immediate early (IE), early (E), and late (L) gene expression. This classification is based on the resistance and susceptibility to chemical inhibitors of protein synthesis (Cycloheximide) and viral DNA replication (e.g., phosphonoacetic acid [PAA] or Ganciclovir [GCV]). The viral IE2 protein serves as an essential transactivator and is required for the transcription of early genes. Late gene expression is dependent on viral genome replication and the activity of the viral late transcription factor complex (LTF) ^6,7^. LTF binds to TATT sequences in viral promoters instead of canonical TATA boxes ^8–10^. In contrast to this functional classification, recent mass spectrometry data defined five temporal classes characterized by distinct protein expression patterns ^11^. Precision nuclear run-on of capped RNA fragments (PRO-cap) experiments recently identified 7,478 viral transcription start site regions (TSR) upon infection of primary fibroblasts ^12^, attributable at least in parts to the activities of IE2 ^13^ and/or LTF ^7^. The complexity of gene expression of the 235 kb viral genome thereby drastically overshoots even the most optimistic estimates. The molecular mechanisms governing their biogenesis and kinetic regulation remain poorly understood.

Here, we provide a detailed and time-resolved map of transcriptional activity of the HCMV genome revealing >2,600 stable HCMV-encoded transcripts. Integrative analysis of previously published PRO-seq, Ribo-seq, and proteomics data revealed that the more complex temporal kinetics of HCMV protein expression ^11^ are governed by (*i*) translation of incoming virion-associated viral mRNAs, (*ii*) combined transcriptional output of multiple TSS expressed with distinct kinetics per viral ORF, (*iii*) activation of L gene promoters before the onset of genome replication and subsequent signal amplification, and (*iv*) differences in viral RNA and protein stability. Most importantly, we identify extensive pervasive transcription of the HCMV genome that does not result in stable viral mRNAs and does not contribute to the viral translatome. In summary, our study provides a unifying model of HCMV gene expression that explains many of the surprising results of previous studies and identifies extensive transcription of transient RNAs as a common feature of this large DNA virus with its human host.

## Results

### Establishing bona-fide TSS of stable viral transcripts

To resolve the kinetics of the HCMV transcriptome, we performed transcription start site (TSS) profiling at multiple time points during lytic HCMV infection of primary human fibroblasts using STRIPE-seq ^14^ and dRNA-seq ^15,16^ combined with 1 h of metabolic RNA labeling using 4-thiouridine (4sU) and SLAM-seq to differentiate between RNA levels and *de novo* transcriptional activity (**Fig. 1A** and **Extended Data Fig. 1A**). A systematic analysis of known bona-fide cellular TSS indicated high reproducibility **(Extended Data Fig. 1B**,**C**) and excellent signal-to-noise ratios of >900 for dRNA-seq and >150 for STRIPE-seq (**Extended Data Fig. 1D**). Our GRAND-SLAM analysis pipeline ^17,18^ confirmed the expected T-to-C conversion rates of >2.7% (**Extended Data Fig. 1E-H, Extended Data Tab. 1**), which enabled accurate quantification of newly synthesized and pre-existing RNA for TSS with dozens of reads ^17^. We used our tool iTiSS ^19^ to call viral TSS for each time point and condition. This reproducibly identified 2,606 TSS in both of the two TSS profiling experiments. In line with our previous observations in the HSV-1 genome ^16^ and the human genome ^19^, both dRNA-seq and STRIPE-seq resulted in substantial amounts of putative TSS that were only reproducibly observed by one protocol but not the other (**Fig. 1B, Extended Data Fig. 1A**). 649/2,606 (24.9%) of the TSS observed by both techniques could be validated by low coverage 3^rd^ generation sequencing data ^20^, but merely 214/2,217 (9.65%) and 384/5,633 (6.82%) of the TSS identified only by dRNA-seq or STRIPE-seq, respectively (**Fig. 1B**). While at least some of the viral TSS observed by only one of the two approaches thus presumably represent *bona fide* viral TSS, we decided to focus on the 2,606 high quality TSS. Late in infection, >70% of newly synthesized TSS were viral in both the STRIPE-seq and dRNA-seq data (**Fig 1C**). We integrated these and publicly available HCMV data into a genome browser and a web-based visualization platform (see https://doi.org/10.5281/zenodo.5801030 and https://erhard-lab.de/web-platforms).

**Figure 1.**
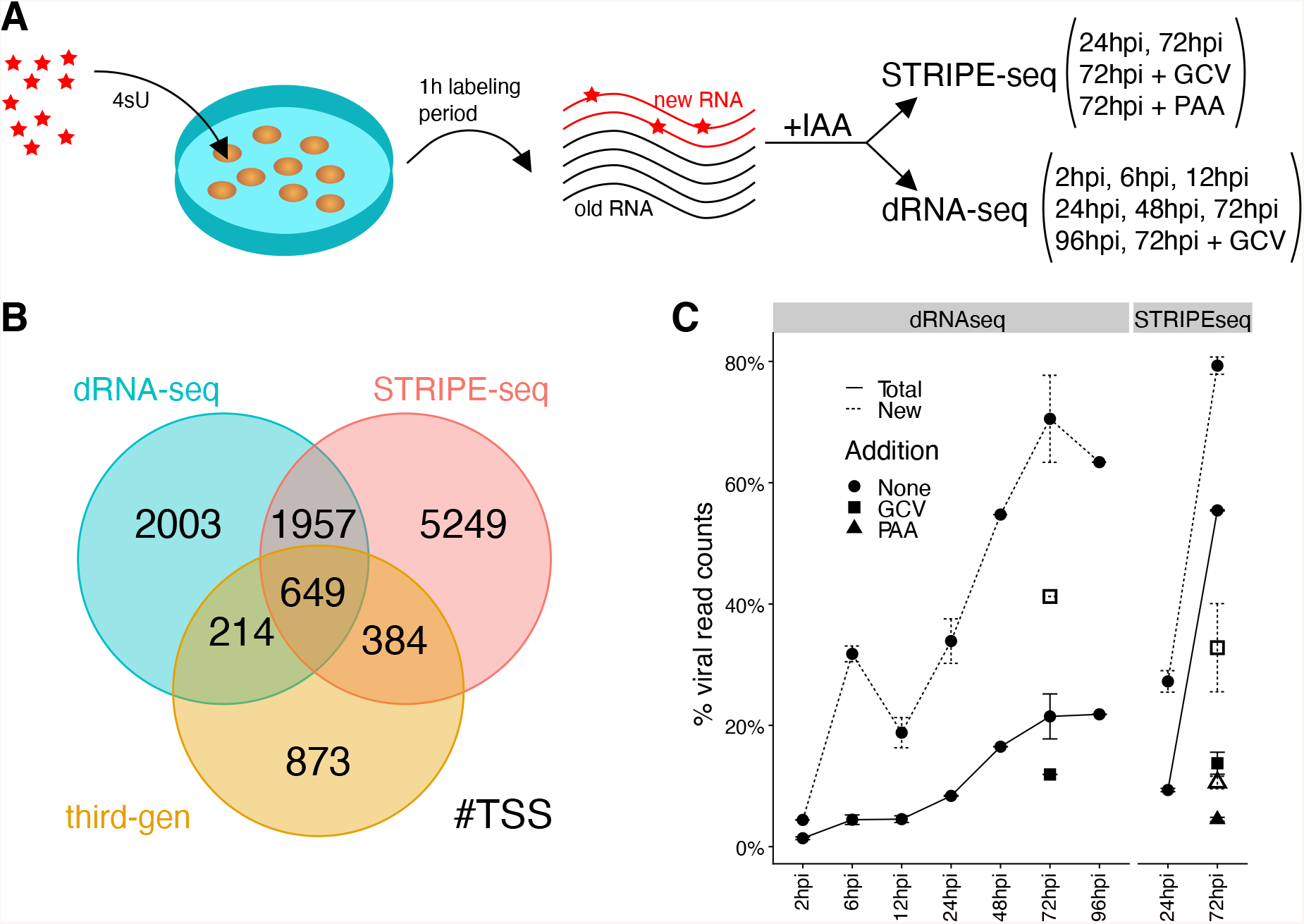
**A**. Schematic overview of the transcription start site profiling experiments. HCMV-/mock-infected cells were exposed to 4-thiouridine (“4sU”) for one hour prior to harvesting RNA at the indicated time points. Sequencing libraries were prepared using STRIPE-seq ^14^ or dRNA-seq ^15^ after converting 4sU into a cytosine analog using iodoacetamide (IAA). The TSS profiling experiments were performed from independent experiments and as two biological replicates. **B**. Venn diagram depicting the overlap of predicted TSS in dRNA-seq, STRIPE-seq and third-generation sequencing approaches using iTiSS ^19^. **C**. Relative amounts of total (solid line) or new (dotted line) viral RNA among overall total or new RNA for dRNA-seq and STRIPE-seq, respectively. Ganciclovir (GCV) and phosphonoacetic acid (PAA) treated samples are indicated (filled rectangle/triangle = GCV/PAA total RNA, rectangle/triangle without fill = GCV/PAA new RNA).

### Distinct promoter sequence motifs govern the kinetics of viral gene expression

Except for TATA and TATT motifs ^8,21^, as well as NF-κB binding sites in the major immediate early promoter (MIEP) ^22,23^, little is known about cis-regulatory motifs driving the expression of viral promoters. To screen for novel sequence motifs, we aligned viral promoter sequences at the corresponding TSS and computed sequence logos for the -100 to +10 nt region focusing on the 500 most strongly expressed TSS in either human or HCMV genome at each time point (**Extended Data Tab. 2**). This revealed a strong pyrimidine/purine (“PyPu”) motif (Inr element) at the viral TSSs ^24^, and a TA-rich element (TATA or TATT boxes) ∼30bp upstream thereof (**Fig. 2A, Extended Data Fig. 2A**,**B**). The Inr element as well as TATA motif are bound by different components of the transcription factor II D (TFIID) complex. The TATT motif is recognized by the viral LTF ^8^. PyPu motifs occurred independently of viral transcriptional activity, were maintained throughout infection, and were significantly more frequent for viral TSS than for cellular TSS (**Fig. 2B-C**, p-value<2.03^-4^; Fisher’s Exact Test). In the host genome, the strongest promoters had less frequent PyPu (p value=1.01^-111^, Fisher’s exact test), which we found to coincide with an alternative cellular initiation mechanism via the TCT motif (**Extended Data Fig. 2C, Extended Data Tab. 3**). TATW boxes occurred more frequently in the promoters of the top 100 most strongly expressed genes (**Fig. 2B**; HCMV: p-value < 10^−3^, cellular: p-value < 10^−5^; Fisher’s Exact Test) in both cellular and viral genomes. The usage of viral TATT box promoters markedly increased already at 24 hpi (**Fig. 2C**). Overrepresentation of cellular TATA-TSS declined after 6 hpi (p-value < 0.012; Fisher’s Exact Test) suggesting that TFIID is sequestered to the viral genomes already in the early phase of infection. TATA boxes are located 25-33 bp upstream of the TSS ^25^. Our data confirmed this for both cellular as well as HCMV genes (**Fig. 2C**). Interestingly, however, the location of viral TATT boxes was shifted 2-3 bp further upstream (**Fig. 2D**, p-value<10^−5^; Fisher’s Exact Test). We concluded that HCMV generally utilizes PyPu-dependent initiation and that HCMV evolution favored strong promoters transcribed via canonical initiation mediated by TFIID or its own LTF complex.

**Figure 2.**
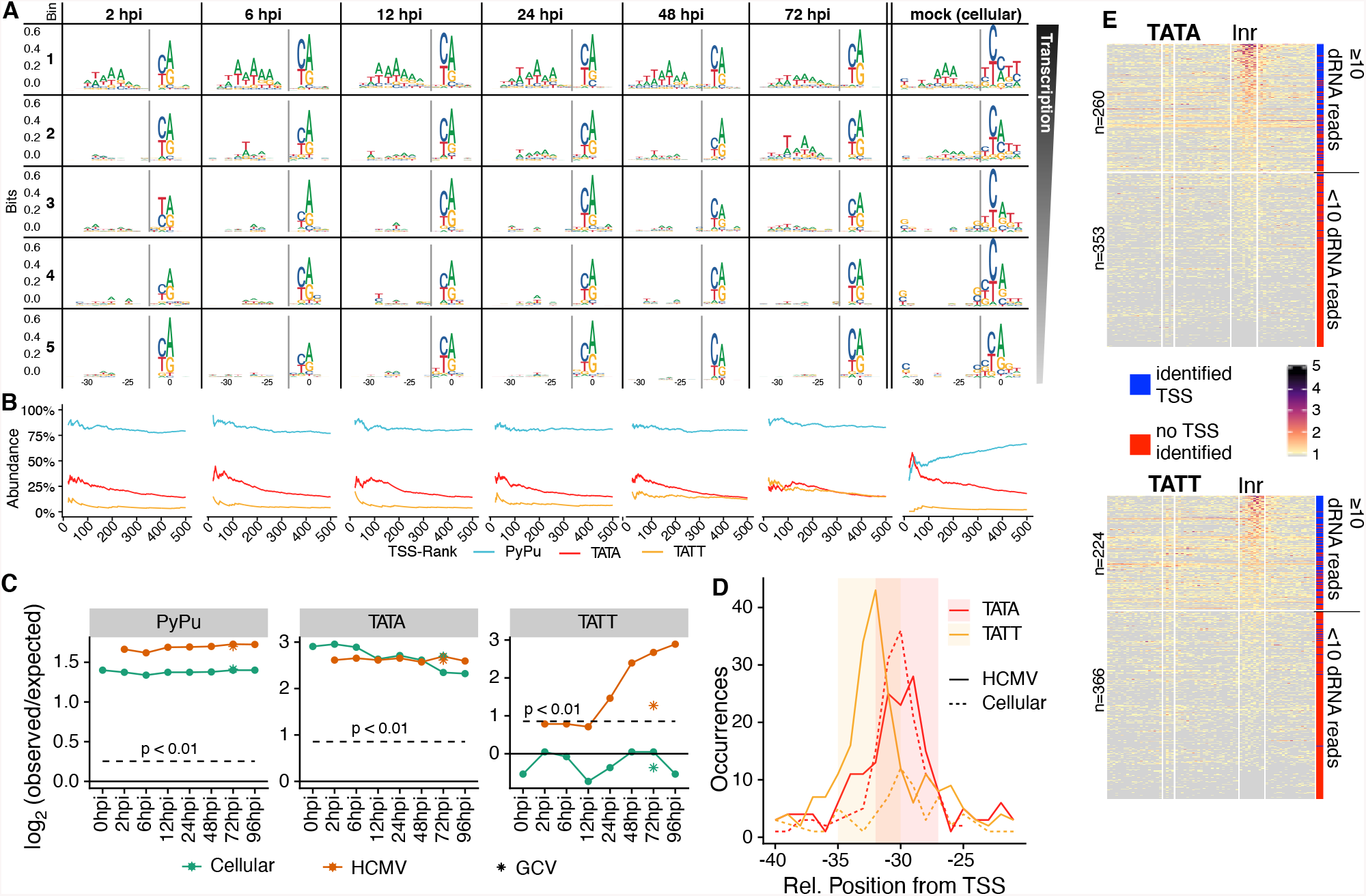
**A**. Sequence logos showing the -32 to -23 bp and the -2 to +2 bp of the 500 most highly expressed TSS during each time point (from left to right) in our dRNA-seq dataset divided into 5 bins according to expression strength (from top to bottom). See also Extended Data Fig. 2A-B for the full +/-100 bp around the TSS. **B**. Line plots showing the percentage of promoters containing correctly positioned (see Extended Data Tab. 3) PyPu, TATA and TATT motifs among the top n TSS on the x-axis. Because of the high variance due to the low numbers of TSS, the first 20 ranks were discarded in the plots. **C**. Log_2_ odds of observed vs. expected occurrences of correctly positioned PyPu, TATA, and TATT motifs in the human and HCMV genome. The critical region (p<0.01, binomial test) is indicated as a dashed line. **D**. Position distribution of TATA (red) and TATT (yellow) motifs relative to the TSS for cellular (dotted line) and HCMV (solid line) TSS. The distance between the motif start and the TSS is shown. **E**. Heatmaps showing the total 5’-read counts over all time points around all TATA-motifs (top) and TATT-motifs (bottom) found in the HCMV genome. The TATW-motifs as well as the expected region of the TSS are indicated. For each row the bar on the right indicates whether a TSS was identified in the respective region by iTiSS (blue) or not (red). The heatmap is sorted according to the number of reads inside the Inr region. The threshold of ≥10 reads within the Inr region is indicated.

We next asked whether the presence of a TATW-box located 24 to 32 (TATA) and 27 to 35 (TATT) nt upstream of an initiator (Inr) element was sufficient to trigger viral transcription by analyzing all occurrences of these motifs (n=970, TATA; n=973, TATT) on both strands of the HCMV genome (**Fig. 2E**). Interestingly, 29% (TATA, 176/613) and 28% (TATT, 165/590) matched to a site in our high-quality set of viral TSS. Moreover, 42% (TATA, 260/613) and 38% (TATT, 224/590) had a least one position exceeding background (≥10 reads) inside the expected Inr location in our data. Thus, a core promoter consisting of a TATW box and a corresponding Inr element is not sufficient for transcription of a stable viral transcript. Motif searches did not identify motifs differentiating effective from ineffective TATW-Inr sites except for a TA-rich extension of 2 bp downstream of few of the effective TATA and TATT sites and two other spurious motifs (**Extended Data Fig. 2D**). Interestingly, AA and AG extensions of TATA, and AA and TA extensions of TATT were strongly enriched among TSS exceeding background (**Extended Data Fig. 2E**, TATAAA, p-value<3.1×10^−7^, TATAAG, p-value<6.8×10^−5^, TATTAA, p-value<1.3×10^−6^, TATTTA, p-value<1.1×10^−11^), but for each extended TATW motif, ≥45% of all TSS were below background. We concluded that elements other than simple sequence motifs around TATW boxes are necessary to define sufficiency of core promoters for viral transcription.

We next used MEME ^26^ to discover novel putative transcription factor binding sites other than core promoter sequences in the HCMV genome. In our control sets of human promoters, we identified the TATA box as well as the TCT promoter and the GC box, all of which are known cellular promoter sequences ^25,27,28^ (**Extended Data Fig. 2F**). However, none of our comprehensive analyses resulted in any viral motif that did not resemble TATW boxes (**Extended Data Fig. 2F**). Searching for known motifs of transcription factors can improve the sensitivity. We therefore evaluated all known transcription factors (TF) for binding sites in viral promoters using TFM-Explorer ^29^. These analyses revealed 9 significantly enriched TF-binding motifs in the top 100 promoters of HCMV at different time points (**Extended Data Tab. 4**). In addition to the TATA-binding protein (36 sites, p-value=8×10^−10^), the most strongly enriched TF was HIF-1α (34 sites, p-value=10^−10^), which has been described previously to be induced in the first few hours of HCMV infection ^30^ and to act as an antiviral host factor ^31^. The enrichment of HIF-1α target sites in viral promoters indicates that HCMV usurped this host factor to regulate the expression of dozens of viral genes. Another top hit in our list was MYC (18 sites, p-value=4×10^−7^), which has been described to be activated by IE1 and IE2 ^32^. Interestingly, MYC target sites were not enriched before 6 hpi. This is consistent with our previous findings from murine CMV ^33^ and indicate that HCMV activates MYC to enhance the expression of viral genes ^34^. In summary, extensive sequence analyses of our TSS data revealed viral transcription initiation to depend uniformly on the PyPu element, that the strength of transcriptional activity throughout the course of infection is predominantly governed by TATW boxes, and that HCMV usurped specific cellular TFs including HIF-1α and MYC to facilitate the expression of specific viral genes.

### LTF drives transcription prior to viral genome replication

TATT motifs are bound by the viral LTF and drive viral late gene expression upon viral genome replication ^35^. However, TATT promoters were also already significantly enriched for the top 500 TSS at 12 hpi and despite blocking viral genome replication (72 hpi+GCV; **Fig. 2C**). This was further confirmed by a metagene analysis of transcriptional activity (**Fig. 3A**), where relatively high read levels were observed at TATT-TSS without genome replication (12 hpi and 72 hpi+GCV) whereas transcriptional activity at 2 hpi did not exceed background signal. For TATA-TSS the 72 hpi+GCV data also showed a metagene profile that was distinct from samples not treated with GCV.

**Figure 3.**
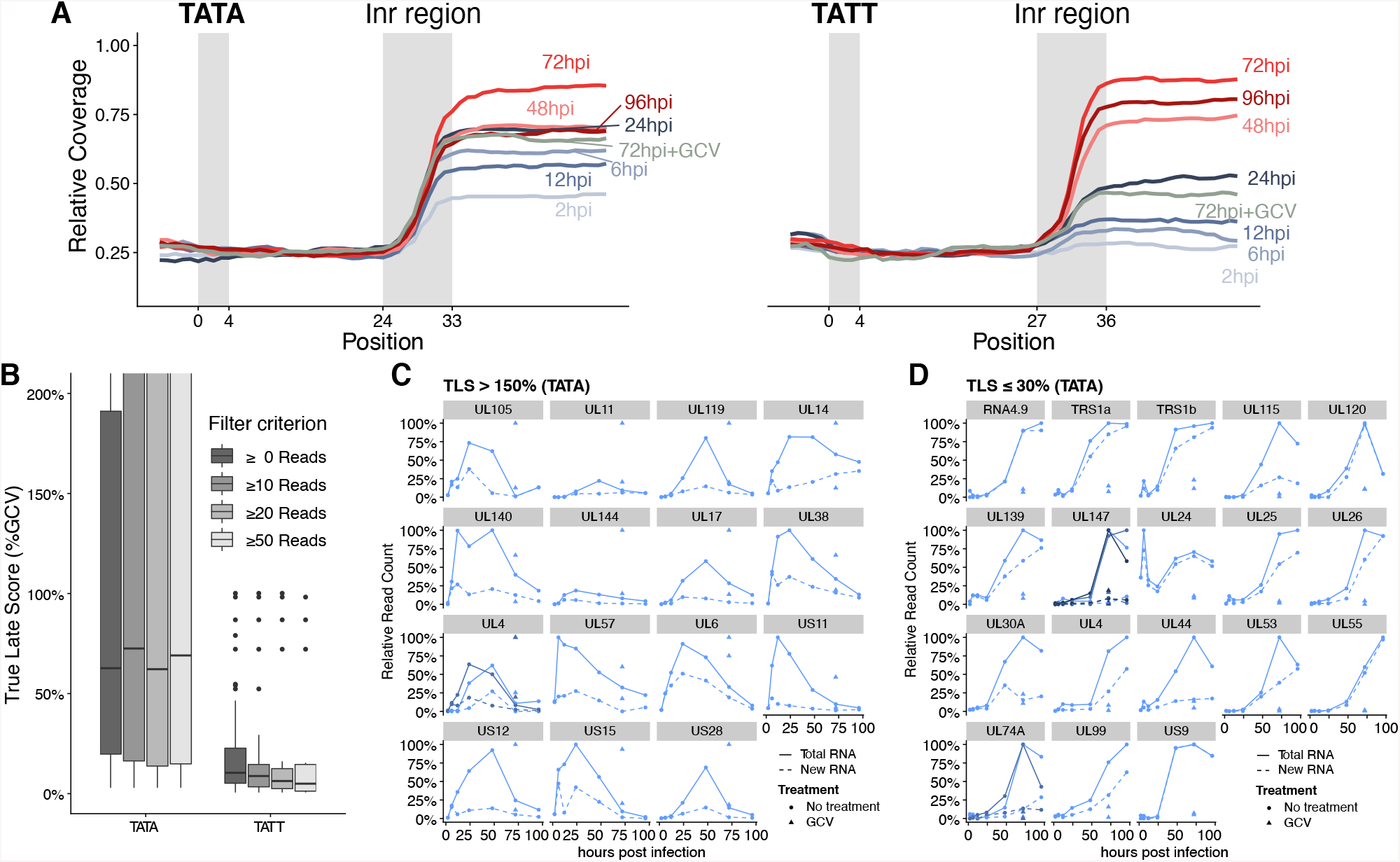
**A**. Metagene plots for each time point centered around all TATA or TATT-motifs with identified TSS in the HCMV genome. Only reads with at least 2 T->C mismatches are considered. The respective TATW-motif as well as the region of the expected TSS are indicated. See Extended Data Fig. 3A for a metagene plot involving all reads. **B**. Boxplots showing the “True Late Score” (TLS; ratio of normalized expression values 72 hpi+GCV vs. 72 hpi) for all TATA- or TATT-TSS, TSS with ≥10 reads, TSS with ≥ 20 reads and TSS with ≥ 50 reads, respectively. **C**,**D**. Abundance profiles for all TATA-TSS with at least 20 reads, TLS>150% (**C**) or TLS<30% (**D**) and a canonical ORF located within 1 kb downstream for total RNA (solid line) and newly synthesized RNA (dashed line). GCV treated samples are indicated by a triangle. For each TSS, the upper triangle depicts total RNA and the lower newly synthesized RNA. If multiple TATA-TSS are present for a single ORF, all TSS are shown and shaded from light blue to dark blue.

To investigate the dependency on genome replication for individual TSS with TATA and TATT boxes, we defined the “true-late score” (TLS) as the percentage of normalized TSS reads in the 72 hpi+GCV compared to the 72 hpi data for TSS (**Fig. 3B**). TATA-TSS showed a surprisingly broad TLS distribution, independent of expression levels. For 33% of the TATA-TSS, GCV treatment substantially increased RNA levels at 72 hpi (TLS>150%, **Fig 3C, Extended Data Fig. 3A**). All these TSS have in common that they are strongly expressed early and downregulated late in a GCV-susceptible manner. Furthermore, 41% of the TATA-TSS had a TLS <30%, i.e. were sensitive to GCV treatment (**Fig. 3D, Extended Data Fig. 3B**). Most of these TATA-boxes were located 30 to 33 bp upstream of the TSS instead of 29 to 31 bp (**Extended Data Fig. 3C**) suggesting that, at least in part, promiscuous LTF binding instead of TFIID drives their expression. By contrast, TATT-TSS predominantly had TLS in between 5% and 25% (25% and 75% percentiles). We obtained similar results when computing the TLS with the 24 hpi sample instead of 72 hpi+GCV (**Extended Data Fig. 3D**). In conclusion, transcription is already initiated at low levels from TATT promoters in the early phase of infection but is strongly augmented upon genome replication.

### Virion-associated RNAs are less efficiently translated than de novo transcribed viral RNAs

HCMV virions deliver so called “virion-associated” RNAs derived from the producer cell into the newly infected cells ^36,37^. To assess their contribution to viral translation, we focused on TSS that were present in total RNA at 2 and 6 hpi but did not show evidence of *de novo* transcription based on U-to-C conversions in our dRNA-seq data. Consistent with the defined role of TATT motifs in viral late gene expression, these characteristics were prevalent for TATT-TSS but not TATA-TSS (**Fig. 4A**). To test whether virion-associated transcripts are productively translated, and to quantify their translational efficiency, we compared our TSS profiling data to Ribo-seq data ^4^. To exclude translational noise from other viral transcripts, we only considered TSS that were identified at 2 hpi and are located at most 1 kb upstream of canonical large ORFs. We found 28 TATA-associated and 17 TATT-associated TSS that passed our stringent filtering. Their translational efficiencies, defined as the translation strength per mRNA of TATA-associated TSS, were significantly higher than those of TATT-associated TSS during the first six hours of infection (**Fig. 4B**, Mann-Whitney-U Test p-value: 6 hpi = 0.03). However, this already reverted by 12 hpi, which was confirmed in the Lactimidomycin (LTM)-treated samples (**Extended Data Fig. 4A**,**B**). We concluded that virion-associated RNA is successfully imported into newly infected cells. However, the respective transcripts are not as efficiently loaded with ribosomes as newly transcribed viral RNAs.

**Figure 4.**
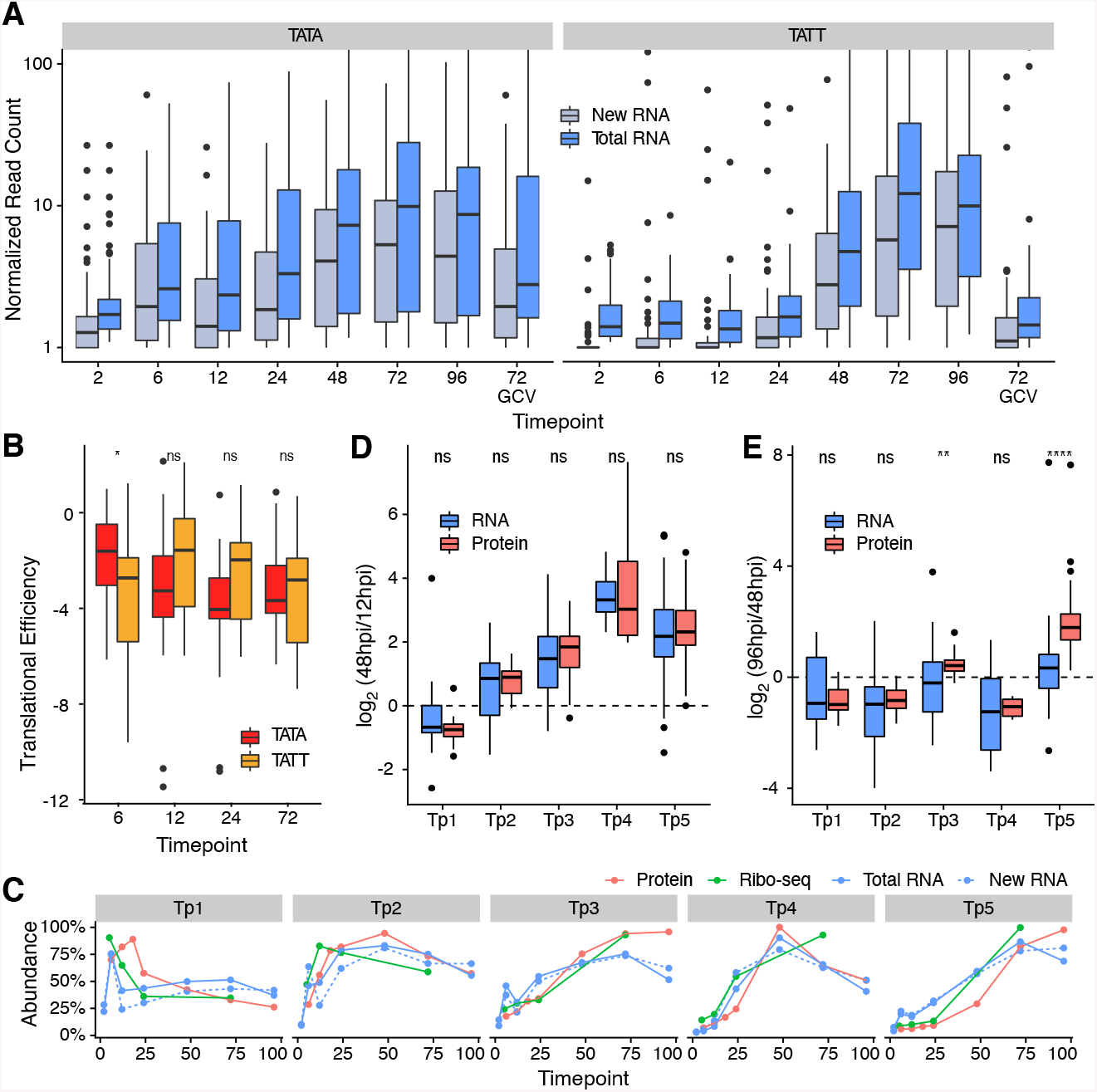
**A**. Boxplots showing the normalized read counts for total (blue) and new (gray) RNA over the course of infection for TATA (left) or TATT (right) associated TSS identified at 2 hpi. **B**. Translational efficiency (translation / transcription) for TATA and TATT associated TSS identified at 2 hpi. Only TSS upstream of canonical large ORFs were included. Translation strength was estimated from the Ribo-seq codon count in the cycloheximide-treated samples (normalized by length). Statistical significance (Mann-Whitney-U test) is indicated (*: p<0.05; the p-values were: 6 hpi, p = 0.033; 12 hpi, p = 0.06, 24 hpi, p = 0.68, 72 hpi, p = 0.82). **C**. For all temporal protein (Tp) clusters defined by Weekes et al. ^11^, the relative protein abundance (red), the translation strength according to Ribo-seq (green), RNA levels according to dRNA-seq (blue, solid lines), and transcriptional activity according to new RNA in dRNA-seq (blue, dashed lines) are shown. All TSS within at most 1 kb upstream of a canonical ORF were considered. **D**. Comparison of the 48 hpi vs. 12 hpi fold changes for TSS expression and protein abundances for each of the five Tp-clusters. For none of the Tp-clusters the difference is significant (Mann-Whitney-U test; p-values: Tp1=0.63, Tp2=0.82, Tp3=0.35, Tp4=0.75, Tp5=0.61). **E**. Comparison of the 96 hpi vs. 48 hpi fold changes for TSS expression and protein abundances for each of the five Tp-clusters. Only for clusters Tp3 and Tp5 the differences were significant (Mann-Whitney-U test; **: p<1.0^-2^, ****: p<1.0^-4^; p-values: Tp1=0.83, Tp2=0.55, Tp3=0.006, Tp4=1.0, Tp5=2.2^-11^).

### Integrative analysis predicts post-translational regulation during HCMV infection

To investigate how IE, E, and L proteins depend on gene regulation exerted at the level of transcription, we performed integrative analyses of genome-wide data on transcriptional activity (TSS profiling, new RNA), mRNA levels (TSS profiling, total RNA), translational activity (Ribo-seq ^4^), and protein levels (mass spectrometry ^11^). We first focused on the five temporal protein classes (Tp1-5) defined by Weekes *et al*. ^11^. Interestingly, their average mRNA profiles closely resembled those of their corresponding proteins with respect to proteomics and Ribo-seq data, except for a drop in RNA levels for Tp3 and Tp5 at 96 hpi that was not accompanied by decreasing protein levels (**Fig. 4C**). Indeed, changes in individual protein levels from 12 hpi to 48 hpi for Tps closely resembled changes in their RNA levels (**Fig. 4D, Extended Data Fig. 4C)**. From 48 hpi to 96 hpi, downregulation of Tp1, Tp2 and Tp4 proteins was also paralleled on the RNA level. However, for Tp3 and Tp5 the increase in protein levels was much more pronounced than for their corresponding RNAs (**Fig. 4E**). This suggests that RNA levels have largely reached steady state levels by 48 hpi, whereas protein levels still increase until 96 hpi for Tp3 and Tp5. We concluded that the mRNA kinetics inferred from our TSS data are consistent with published proteomics data, and that the remaining differences are likely consequences of RNA turn-over being faster than protein turn-over.

### A subset of HCMV proteins is translated from multiple mRNAs with distinct kinetics

For 111 out of 162 canonical ORFs, we identified more than one TSS located 1kb upstream. To analyze their kinetic behavior, we clustered all 516 TSS upstream of canonical ORFs into five temporal RNA classes (Tr1-5; **Extended Data Fig. 4D**). Interestingly, the Tr class of the most abundant (dominant) TSS per ORF only recapitulated the Tp class in 42% of the cases (**Fig. 5A**). Examples such as UL19 indicated that the kinetics of translational activity of an ORF might be a result of the combination of kinetically distinct mRNAs (**Fig. 5B**). Thus, we set up a model to calculate the correlation between translational activity with either the dominant TSS alone or a combination of all TSS (see Methods). From the 63 ORFs that had TSS from multiple Tr classes, 43 ORFs showed significantly improved correlations when considering all TSS (**Fig. 5C**). We conclude that the temporal expression kinetics of dozens of HCMV proteins are governed by distinct transcriptional regulation of multiple TSS.

**Figure 5.**
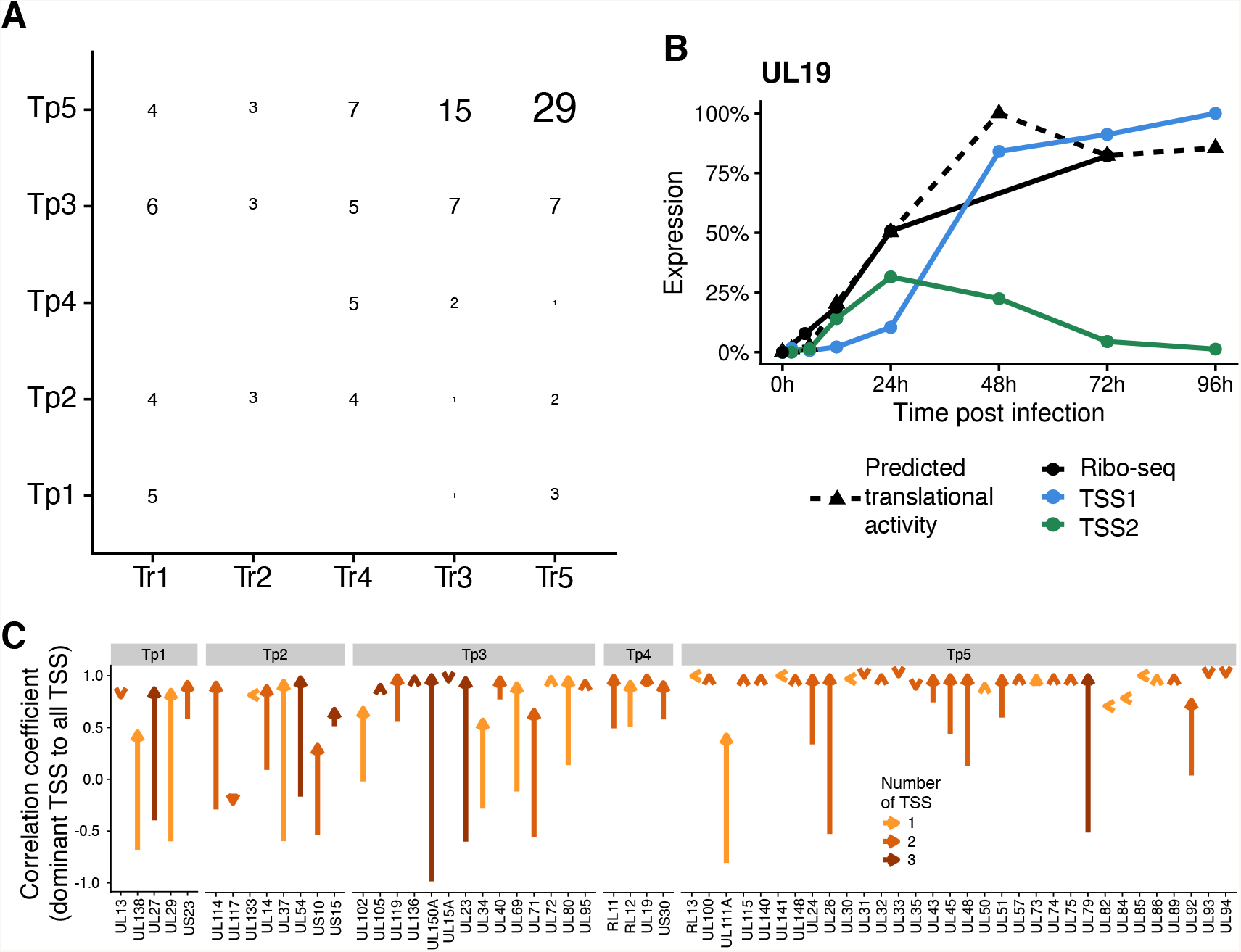
**A**. Comparison of the temporal protein (Tp) classification and the corresponding temporal RNA (Tr) classification of the corresponding dominant TSS (n=117). The numbers indicate the number of ORFs belonging to a specific combination of Tr and Tp class. **B**. Abundance profile of the two TSS for UL19 (blue and green lines). The solid black line depicts translation strength obtained from Ribo-seq data. The dashed black line is the predicted translation using our model (see Methods) using the two TSS. **C**. Improvements of the correlation between the transcription (TSS-profiling data) and translation (Ribo-seq data) by using only the dominant TSS (start of the arrow) or a combination of all TSS (tip of the arrow) for each gene with >1 TSS. The colors show the number of available distinct TSS by their Tr class.

### Pervasive non-productive transcription in HCMV infection

Recently, Parida *et al*. ^12^ utilized PRO-seq and PRO-cap experiments to identify over 7,000 transcription start site regions (TSRs) active at 96 hpi in the HCMV genome. We mapped the previously identified PRO-cap TSRs to the TB40E-Lisa reference genome, which left us with 6,985 TSS (see Methods). Interestingly, we recovered only 946 of these at 96 hpi in our TSS data, and 1,712 when pooling all data. Heatmap analysis showed that this was not simply a consequence of lower sensitivity due to differing read depths between these data sets, as the distribution of PRO-cap read counts for TSS detected in our data resembled the read count distribution of PRO-cap-only TSS (**Fig. 6A**). Furthermore, we reproducibly identified >1,100 TSS that had not been detected by PRO-cap. Importantly, these TSS nevertheless showed the characteristic TATW-boxes and PyPu motifs confirming that they indeed represent *bona fide* TSS (**Extended Data Fig. 5A**), suggesting that they were either no longer transcribed at 96 hpi or not detectable by PRO-cap.

**Figure 6.**
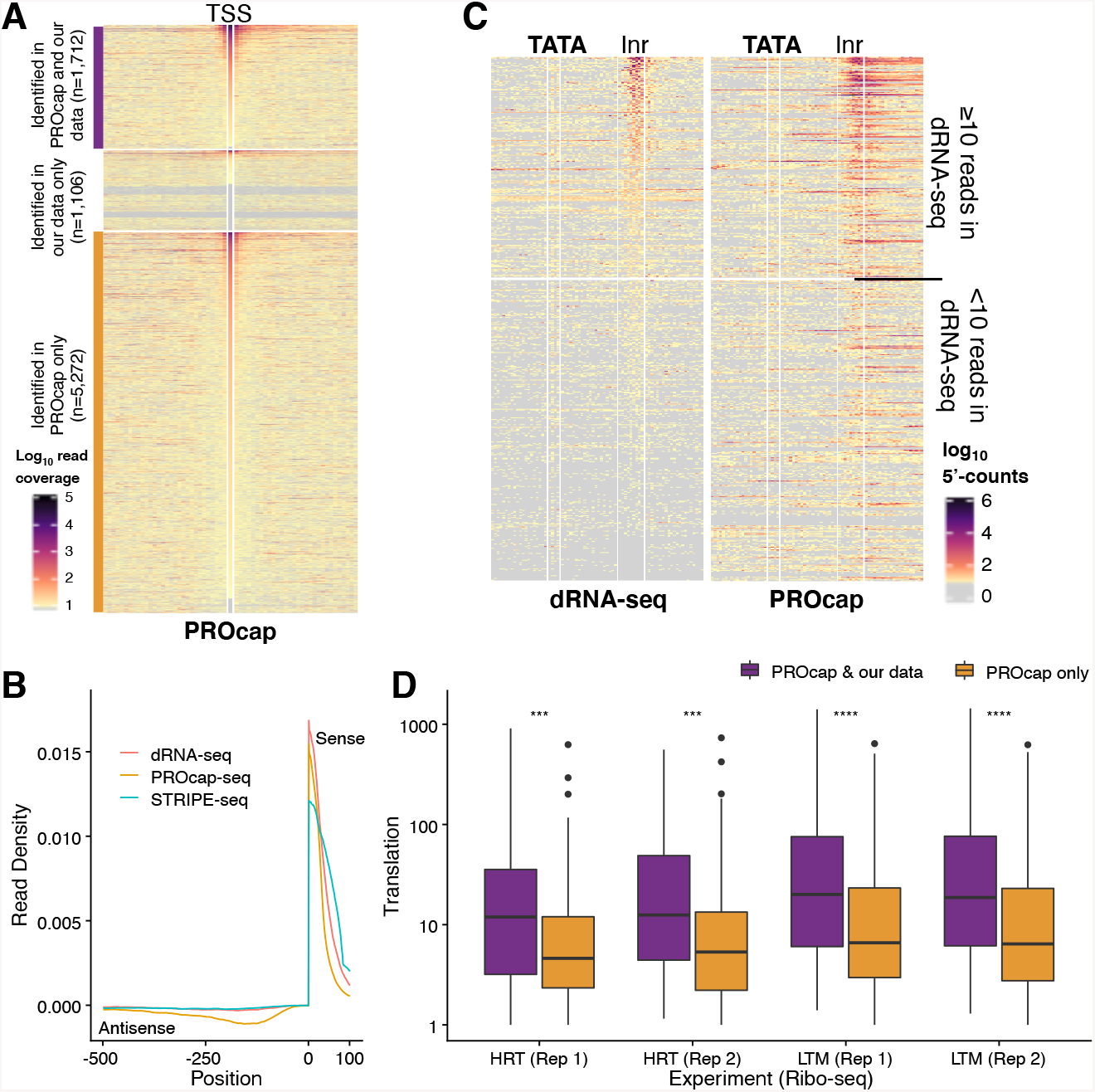
**A**. Heatmaps of the read densities for +/-30bp around TSS at 96 hpi in the PRO-cap data. The upper part shows TSS identified in both PRO-cap and dRNA-seq data, the middle part TSS identified in dRNA-seq only, and the lower part TSS only identified in the PRO-cap data. The rows in each part are sorted by read counts in the PROcap data. **B**. Metagene plot for dRNA-seq, PRO-cap and STRIPE-seq of cellular TSS found at annotated protein-coding transcripts. Downstream of the TSS read counts are shown for the sense strand and upstream of the TSS the read counts are shown for the antisense strand. **C**. Heatmaps showing the total 5’-read counts over all time points in the dRNA-seq data (left) and the total 5’-read counts at 96 hpi in the PROcap data (right) around all TATA-motifs found in the HCMV genome. The TATA-motifs as well as the expected region of the TSS are indicated. The heatmap is sorted according to the number of reads inside the Inr-region of the dRNA-seq data. The threshold of ≥10 reads within the Inr region in the dRNA-seq dataset is indicated. **D**. Comparison of the translation rates downstream of TSS identified in both data sets (PRO-cap and dRNA-seq) or only in PRO-cap inferred from the harringtonine (HRT) or lactimidomicin (LTM) treated Ribo-seq samples. PRO-cap only TSS were selected to match the expression strength of TSS identified in both data sets. The differences were significant (Mann-Whitney-U test; ***: p<1.0×10^−3^, ****: p<1.0×10^−4^; p-values: HRT (Rep 1)=1.2×10^−4^, HRT (Rep 2)=1.2×10^−4^, LTM (Rep 1)=6.5×10^−6^, LTM (Rep 2)=5.7×10^−6^)

It is important to note that both dRNA-seq and STRIPE-seq identify TSS of stable viral transcripts that accumulate during infection whereas PRO-cap measures nascent transcription initiation at a define time of infection. Indeed, metagene analysis in the human genome identified thousands of PROMPTs ^38^ in the PRO-cap but not in the dRNA-seq or STRIPE-seq data (**Fig. 6B**). PROMPTs correspond to transcription initiation events antisense upstream to TSS of protein-coding genes that give rise to transient RNAs. We thus asked whether PRO-cap-only TSS might represent a much larger transient HCMV transcriptome. We first assessed how many TATW-motifs associated with Inr elements correspond to PRO-cap-only TSS. 208 out of 613 TATA-Inr and 214 out of 590 TATT-Inr promoters had at least 10 reads in the PROcap data but were not detectable by dRNA-seq (**Fig. 6C** and **Extended Data Fig. 5B**). Taken together, our data indicate that 40% of all TATW-Inr promoters initiate transcription of stable RNAs, and an additional 30% result in transient RNAs. While TATW-Inr elements in the viral genome are thus sufficient to initiate transcription in >70% of cases, additional signals appear to be required to promote efficient transcription initiation as well as efficient elongation of viral transcripts.

Next, we utilized Ribo-seq data to analyze translation initiation at the first AUG start codon downstream of individual TSS. PRO-cap-only TSS showed significantly weaker translation initiation rates downstream compared to TSSs from stable viral transcripts for equally well transcribed TSS in the PRO-cap data (p-value<1.3×10^−4^, Mann-Whitney-U-Test, **Fig. 6D** and **Extended Data Fig. 5C**,**D**). Thus, Ribo-seq data provides independent evidence that PRO-cap-only TSS represent unstable RNAs. Moreover, we found that 3,268 of the 5,272 (62%) PRO-cap-only TSS were located within 100-350 bp antisense upstream of a TSS from our reproducible set of stable TSS (**Extended Data Fig. 5E**). Reminiscent to cellular genes, these represent candidates of viral PROMPTs. Finally, we hypothesized that at least a small fraction of PRO-cap-only TSS should have been captured by trace amounts of dRNA/STRIPE-seq reads. For strong (≥1,000 reads) PRO-cap-only TSS, we found 221 reads in our dRNA-seq and 45 reads in our STRIPE-seq data. In regions 25 bp upstream of PRO-cap-only TSS that we included as control, we only found a single read in dRNA-seq and six STRIPE-seq reads. At PRO-cap-only sites, 33.6% of the dRNA-seq reads had at least one U-to-C conversion. Considering read lengths and U-to-C conversion rates, this corresponded to a new-to-total RNA ratio (NTR) of 0.57 to 1.0 with a mean NTR of 0.78. We concluded that a large number of PRO-cap-only TSS represent *bona-fide* sites of transcription initiation that give rise to short-lived viral RNAs that do not contribute to the viral translatome.

## Discussion

Recent studies identified >7,000 sites of transcription initiation in the 235 kb genome of HCMV ^12^ and discovered translation of >1,000 viral ORFs ^4,5^. Here, we provide a unifying model of the kinetic regulation of this promiscuous HCMV gene expression during lytic infection of primary human fibroblasts. Deep transcription start site profiling based on total cellular RNA combined with metabolic RNA labeling ^18,39^ at various times after infection identified a reproducible set of 2,668 stable viral transcripts. By 72 hpi, >70% of transcription initiation was viral highlighting how efficiently the virus takes over its host cell’s transcriptional machinery. While PyPu elements defined the site of transcription initiation, TATW motifs were associated with strong transcriptional activity. TATA boxes occurred throughout infection in strong promoters, whereas TATT promoters were found for strongly expressed viral late genes. Interestingly, TATA and TATT promoters exhibited distinct sequence motifs directly following TATW, and their position was on average shifted by 2 bp further upstream of the TSS. This indicates that the LTF, which has been shown to bind TATT sequences ^35^, is structurally distinct from the TATA-binding TFIID.

Overall, 1,331 of the 2,668 TSS we identified were located within 1 kp upstream of one of the 168 canonical ORFs. Interestingly, the kinetics of the dominant TSS only matched protein kinetics for 70 of the 162 canonical viral ORFs. For 41 viral ORFs, protein expression kinetics could only be explained by combining translation from more than one transcript isoform. Expression of the respective viral proteins is thus governed by multiple TSS with distinct kinetics. This does not only imply two independent promoters that exhibit specific regulation over time, but also the potential of additional regulation at translational level by the hundreds of viral upstream open reading frames (uORFs) ^4,5^.

In addition to measuring transcriptional activity instead of RNA levels, our metabolic labelling approach enabled us to accurately quantify virion-associated RNA. RNA incoming with virus particles has previously been shown to be translated ^37^. However, whether virion-associated RNA is as efficiently translated as *de novo* transcribed viral RNA remained unclear. Efficient translation depends on effective formation of polysomes that in turn depend on a number of RNA-binding proteins including the cap-binding protein and poly(A)-binding proteins that are partly loaded onto the mRNA during processing ^40^. Whether or not the incoming RNAs efficiently enter polysomes is unclear. Integrative analysis with Ribo-seq data suggested that the translational efficiency of virion-associated RNA is indeed 2-8x lower than for *de novo* transcribed mRNA in the same sample, arguing against efficient loading onto ribosomes.

Recently published PROcap data suggested over 7,000 TSRs in the HCMV genome ^12^, i.e. a site of transcription initiation every 65 nucleotides on average on both strands of the 235 kb genome. For HSV-1, we could recently show that the vast majority of novel viral ORFs identified by Ribo-seq can be explained by novel viral transcripts that initiated in close proximity upstream ^16^. Accordingly, if all of these >7,000 TSRs would give rise to stable viral mRNAs, the viral translatome would have to be substantially larger and contain a plethora of truncated isoforms of the canonical viral ORFs. Our two TSS profiling approaches on total RNA, however, only confirmed 1,712 TSS of stable viral transcripts detectable in total cellular RNA. Integrative analysis of the available big data and exploiting additional information by metabolic RNA labeling provided strong evidence that the majority of the PRO-cap TSR represent non-productive transcription that gives rise to transient viral mRNAs and thus does not contribute to the viral translatome.

A minuscule percentage of TATA sequences in the human genome represents an actual promoter that drives expression as many of them are located in regions of densely packed chromatin. In the HCMV genome, the situation is different. While the viral genome is at least partially chromatinized, it is less compact than the human genome throughout productive infection ^41^. Interestingly, only about 40% of all >1,200 TATW motifs in the HCMV genome with a properly positioned downstream PyPu element gave rise to stable viral RNAs. However, an additional 30% showed nascent transcription initiation by PRO-Cap. Conclusively, most of the HCMV genome seems to be accessible for the transcriptional machinery. Within the host genome, the factors that differentiate productive from non-productive transcription initiation remain poorly understood but involve positive elongation factors like c-myc ^42^ and splicing ^43^. While cellular factors may also govern productive HCMV gene expression, it is likely that at least one viral factor also plays a role. A recent functional genomics screen revealed perturbation of only two viral genes, namely UL69 and UL112, to create distinct, abortive trajectories ^44^. UL112 has recently been shown to undergo liquid-liquid phase separation to form viral replication compartments in the nucleus ^45^. It may thus not only be essential for viral genome replication but also facilitate recruitment of the transcriptional machinery to the replicating viral genomes. UL69 is a conserved herpesviral transactivator ^46^ which may facilitate productive transcription by its interaction with nascent viral RNAs. Its homologs ICP27 (HSV-1) and ORF57 (KSHV) directly affect the fate of viral mRNAs by facilitating proper termination ^47^ and protecting them from nuclear RNA decay pathways ^48^. It is thus highly likely that pervasive transcription and regulation of transient vs. stable viral mRNAs is not unique to HCMV but typical for all herpesviruses. Viral manipulation of pervasive transcription by specific viral factors may not only govern the temporal kinetics during lytic infection but provide programmed roadblocks that prevent spurious viral protein expression and thus avoid immune recognition during latency and facilitate rapid virus reactivation.

## Methods

### Cell culture, viruses, and infections

Primary human foreskin fibroblasts (HFFs) were cultured in DMEM (Dulbecco’s Modified Eagle Medium) supplemented with 10% [v/v] FBS (fetal bovine serum). MRC-5 fibroblasts were seeded on six-well plates and infected with HCMV strains – TB40E-Lisa and BAC2 ^49^ (AD169VarL) at 10 PFU/cell. 4sU labelling at 500 µM was conducted 1 hour prior to lysis at respective time points. Treatment with GCV (ganciclovir) and PAA (phosphonoacetic acid) was performed at 12 hpi using 50 µM and 250 µg/mL respectively.

### RNA extraction

Cells were lysed in 1mL Trizol and RNA extraction was performed with Zymo research Direct-Zol™ RNA Miniprep Plus kit as per manufacturer’s instructions. Prior to TSS profiling, 4sU labelled RNA was subject to alkylation (SLAM) as previously described ^28^. Briefly, RNA was treated with 10mM Iodoacetamide-50% DMSO solution at 50° C for 15 minutes in 50mM PBS (phosphate buffered saline). The reaction was quenched using excess DTT. RNA was purified using Qiagen RNeasy® Minelute® Cleanup kit and eluted in nuclease-free water.

### TSS profiling

TSS profiling was conducted using dRNA-Seq and STRIPE-Seq on independent samples. dRNA-Seq was performed as described previously ^26^ with minor modifications introduced by the Core Unit Systems Medicine (Würzburg) ^12^. Briefly, isolated RNA was subject to 3’ dephosphorylation prior to enzymatic treatment with Xrn-1 nuclease, which led to strong enrichment of 5’ capped RNA. De-capping was conducted to mediate adaptor ligation using RppH (NEB) followed by library preparation using NEBNext® Multiplex Small RNA Library Prep (Illumina). Pair-ended 2×75 bp sequencing was performed on the NextSeq 500 (Illumina) at Core Unit Systems Medicine. Data analysis revealed contamination of all virus-infected samples with Mycoplasma despite regular PCR testing for Mycoplasma, which resulted from a contaminated HCMV stock.

STRIPE-seq libraries were prepared from 160 ng of iodoacetamide-treated RNA according to the protocol described in Policastro *et al* ^14^. Briefly, RNA was subjected to terminator exonuclease treatment, template switching reverse transcription followed by PCR amplification, and size selection. Libraries were sequenced (100 bp paired-end) on the DNBSEQ-G400 platform at BGI Tech Solutions, Hong Kong. No mycoplasma RNA was detectable in these samples.

### Data analysis, statistics, and reproducibility

All scripts and source codes are deposited at Zenodo to provide complete details about the exact parameters used for the individual programs as well as enabling the full reproduction of all our analysis starting from raw data (https://doi.org/10.5281/zenodo.5801030).

For second generation transcriptomic sequencing data, adapter sequences were trimmed using Trimmomatic ^50^ (v. 0.39). Resulting reads with a length <18 were discarded. The remaining reads were mapped to the ribosomal RNA of homo sapiens using bowtie2 ^51^ (v. 2.3.0) with standard parameters. Only the unmappable reads were subsequently mapped against the combined reference genome of homo sapiens (Ensembl hg38 version 90) and HCMV (strain TB40E-Lisa, GenBank accession number KF297339.1; strain Towne for PROcap only, GenBank accession number FJ616285.1) using STAR ^52^ in paired-end mode. Samtools ^53^ (v. 1.9) was used to sort and index the final BAM-file. For subsequent analysis, the gedi toolkit (https://github.com/erhard-lab/gedi) was used. The BAM-files were converted into CIT-files, which can combine multiple BAM-files into a single file, considerably saving memory and access times. STRIPE-seq data as well as PROcap-seq data contain unique molecular identifiers (UMIs). These were accounted for and removed using the same mismatch correction algorithm as UMItools ^54^. For STRIPE-seq, all reads having a 5’-softclip of length >3 were removed ^14^. Further, to reduce error-rates in STRIPE-seq, we only considered reads with at least one additional duplicate. In case of mismatching nucleotides between duplicates, a majority-vote was used to determine the correct one.

Third generation sequencing reads were trimmed and oriented based on the presence of the 5’-adapter and poly(A) tail. The remaining reads were mapped using GMAP ^55^. The consecutive indexing, sorting and conversion into CIT-files was identical to second generation sequencing data as described above.

For the Ribo-seq data, adapters were trimmed using reaper (kraken v. 15.065 toolkit ^56^). Bowtie ^57^ (v. 1.2) was used to first align against the homo sapiens rRNA genome and secondly, remaining unaligned reads against the homo sapiens (Ensembl hg38 version 90) genome. Sorting, indexing and conversion into CIT-files remains the same as for the aforementioned sequencing data.

For the calculation of the enrichment at TSS, reads +/1 bp around the TSS were collected as well as reads in the 100 bp up- and downstream window. A read was considered in one of these three windows, if and only if its 5’-end was located inside of it. Read counts of the respective windows were normalized and the down- and upstream enrichment was calculated by dividing the TSS-read count by the up- or downstream window read count, respectively. The final enrichment value for each dataset is derived by the median of all the TSS enrichment values.

Correlation between datasets was calculated by extracting the number of reads per bp in the HCMV genome. The read counts per bp were normalized by the total amount of mappable reads per dataset and multiplied by 1,000,000. Finally, the Pearson correlation coefficient of the log_10_ transformed read counts per bp between replicates was calculated. Positions with 0 reads were discarded.

Uridine to cytosine (U-to-C) conversion rates as well as error rates and new to total RNA rates (NTRs) for TSS were estimated using GRAND-SLAM ^17^. Only reads with 5’ ends inside a predicted TSS were considered. To decrease error rates, only the overlapping parts of read pairs were considered for the dRNA-seq sample. For STRIPE-seq, all positions were used as they are already error-corrected during the deduplication process (see above).

Transcription start sites (TSS) were called using iTiSS ^19^ (v. 1.3) with the SPARSE_PEAK module. Reads of the individual replicates per time point were pooled. Subsequently, for each dataset, TSS were merged with a +/- 10 bp windows using the sub-program TiSSMerger2 of iTiSS. Finally, only TSS validated by both (STRIPE-seq and dRNA-seq) datasets were considered high quality TSS. Since early time points for our STRIPE-seq sample were not available, we additionally included dRNA-seq TSS from 2 and 6 hpi, if they were validated by at least 3 out of 4 samples.

Sequence logos were generated using the R-package ‘ggseqlogo’ ^58^. TSS per time point were sorted based on their total read count or newly synthesized read count (*NTR * totalRNA*) and in descending order, respectively. For the prediction of promoter sequences, MEME ^26^ (v. 5.3.3) was used with *-minw 4, -maxw 15*, -*objfun de*, and *-mod zoops* parameters. We used multiple different window sizes around the TSS (−50 to 10, -100 to 10, -200 to 10, -10 to 100, -100 to 100) as well as running MEME only on the top 100 and top 500 most strongly expressed TSS. Furthermore, we performed these analyses also after excluding TSS that were in close proximity to other TSS, which might mislead the motif discovery algorithm.

For each TSS we searched for position specific promoters. The promoters as well as their respective sequences and windows relative to the TSS are as follows (of note: the TSS has position 0): TATA-box (Sequence: TATA, Window: -32 to -24), TATT-box (Sequence: TATT, Window: -35 to -27), PyPu (Sequence: YR, Window: -1 to 0), Inr extended (Sequence: BBCABW, Window: -3, 3), BRE upstream (Sequence: SSRCGCC, Window: -38 to -32), BRE downstream (Sequence: RTDKKKK, Window: -24 to -17), DRE (Sequence: WATCGATW, Window: -100 to -1), TCT (Sequence: YYCTTTYY, Window: -2 to 6). The occurrence per TSS-rank plot (Fig. 2B) was generated by sorting the top 500 TSS in descending order based on their newly synthesized read count per time point. Then, in descending order the fraction of observed PyPu, TATA or TATT, respectively, up to the current rank is calculated and plotted. Because of the high variance due to the low number of TSS, the first 20 ranks were discarded in the plots. To test the differences in PyPu occurrence frequencies between HCMV and cellular promoters, we compared the top 500 viral TSS for each time point against the top 500 cellular TSS in mock. For each comparison Fisher’s Exact test was used with a=number of viral TSS with PyPu, b=number of viral TSS without PyPu, c=number of cellular TSS with PyPu, d=number of cellular TSS without PyPu. For the comparison of PyPu and TATW-box occurrences between strongly expressed TSS and weakly expressed TSS, we binned the TSS for each time point based on the expression strength into 5 bins, each containing 100 TSS. Then, we compared the first bin (top 100 most strongly expressed TSS) with the 5^th^ bin by using Fisher’s Exact Test with a=number of TSS with PyPu or TATW, respectively, in bin 1, b= number of TSS without PyPu or TATW, respectively, in bin 1, c= number of TSS with PyPu or TATW, respectively, in bin 2, d= number of TSS without PyPu or TATW, respectively, in bin 2. To test the presence of a shift of location of the TATT box compared to the TATA-box in virus, we used Fisher’s Exact Test with a=number of TATA boxes starting between 33 and 35 bp upstream of the TSS, b=number of TATA boxes starting between 27 and 29 bp upstream of the TSS, c=number of TATT boxes starting between 33 and 35 bp upstream of the TSS, d=number of TATT boxes starting between 27 and 29 bp upstream of the TSS. To test the decrease in TATA occurrences in the human genome over the course of infection, Fisher’s Exact Test was used with a=number of TSS with TATA in mock, b=number of TSS without TATA in mock, c=number of TSS with TATA at 96 hpi, d=number of TSS without TATA at 96 hpi.

For the generation of the TATW heatmaps, we used the tool *MetagenePlot* (https://github.com/erhard-lab/MetagenePlot). TATW boxes were centered based on their first ‘T’. The 5’-counts of all reads over all time points were pooled and are depicted in the heatmap. A TATW box was considered to be in front of a TSS if a TSS was found 24 to 32 bp downstream of it (starting from the first ‘T’) for TATA and 27 to 35 bp downstream for TATT, respectively.

The True Late Score (TLS) was calculated for each TSS by dividing the sum of the normalized read counts of both replicates at 72 hpi for the sample treated with GCV by the sum of the normalized read counts of both replicates at 72 hpi for the sample treated without GCV.

Metagene plots were also generated by using the tool *MetagenePlot*. Here, we used all the TATW boxes that had a TSS identified downstream (see above). The tool calculates the read coverage for each position. Then, each row (TATW-box and down- and upstream region) is normalized by the total read count. Subsequently, for each position in the metagene analysis, the mean-normalized read coverage is plotted. Further, the metagene plots of all time points were combined by normalizing them such that the mean read coverage value of the region between the TATW-box and the supposed TSS-region is fixed at 0.25.

PRICE ^5^ (v. 1.0.3b) was used to estimate the codon counts (number of reads assigned to a specific codon) indicating the positions of the ribosome during translation. The codon counts per codon were normalized by the total codon count times 1,000,000.

Translational efficiencies of virion-associated RNA were calculated once by obtaining the codon count throughout the coding sequence of the respective ORF for our cycloheximide-treated samples and additionally normalizing it by the ORF’s length. For our samples treated with lactimidomycin, we only obtained the codon count at the respective ORF’s start codon. Further, in both cases, the obtained codon count is divided by the read count at the corresponding TSS. Only ORFs, where the most strongly transcribed TSS in a 1kb upstream window at any time point was either TATA- or TATT-associated were considered.

For each TSS, the next protein-coding downstream ORF found in a 1kb window and annotated in HCMV strain TB40E-Lisa (GenBank accession number KF297339.1) was considered to be its respective transcribed ORF. If no ORF was found, the TSS was not assigned any ORF. For each ORF, the assigned TSS with the maximal-normalized read count at any time point was considered its dominant TSS. Any TSS, where the normalized read count during any time point did not exceed 10% of the dominant TSS for the respective ORFs were discarded.

Protein abundances were obtained from Table S2 of Weekes *et al*. ^11^ and their associated temporal classes from Table S6C. Table S6C were missing some temporal classifications, which we manually added based on their Data S1 plots. Comparison of transcriptional expression and the protein abundances was performed for all proteins, where at least one TSS was found. Per protein, the read counts over all time points were normalized by the maximum read count. Then, the mean read count per time point over all TSS of the respective temporal class (taken from the associated protein) were plotted.

The transcription classes (Tr) were assigned to the TSS by first clustering the dominant TSS for each ORF and subsequently using the resulting centromeres to cluster the remaining TSS. Only TSS, which were expressed with at least 10% of the expression strength of the main TSS per ORF were used. Expression strength of an individual TSS is depicted by the timepoint with the highest normalized read count. For each dominant TSS, the read counts of all time points were normalized by dividing them by the maximum read count of any time point for the respective TSS (i.e. the time point with the highest read count is subsequently 1). Then, the KMeans++ clustering algorithm ^59^ was used on the data from 6, 12, 48, 72 and 96 hpi with k=5 to cluster the TSS into 5 transcriptional classes. The clustering was performed multiple times, always resulting in similar profiles. Consequently, in order to assure a deterministic clustering to ease subsequent automatic analysis steps, we set the seed to 42 for the KMeans++ algorithm.

To compute the correlation of the temporal Ribo-seq profile with the combined TSS profiles, we used non-linear least squares regression (nnls package version 1.4) to fit a model predicting the overall translational activity of an ORF based on the translational efficiency of each of its TSS. In this model, the Ribo-seq signal of the main ORF *M*_*j*_ in sample *j* is related to the dRNA-seq signal *u*_*i,j*_ of TSS *i* in sample *j* and the translation rate *R*_*i*_ of the main ORF from TSS *i* via *M*_*j*_ *=* ∑_*i*_ *u*_*i,j*_ *R*_*i*_. When there are more samples than TSS, this is an overdetermined system of linear equations and the non-negative coefficients *R*_*i*_ can be estimated by regression. The correlation was then computed for *M*_*j*_ vs. ∑_*i*_ *u*_*i,j*_ *R*_*i*_.

TSS were predicted as described by Parida *et al*. ^12^, by first aligning the PRO-cap data to the Towne strain of HCMV (GenBank accession number FJ616285.1), removing duplicates using the provided UMIs and then running their tool TSRFinder on it. As described ^12^, we merged consecutive TSRs and took the position with the highest read count as the TSS. To make these TSS comparable to our data, which were mapped to TB40E-Lisa, we extracted the sequences +/-40 bp around the TSS from the Towne sequence and mapped these to the TB40E-Lisa sequence using STAR with the “*--alignEndsType EndToEnd*” option.

For the comparison of translation rates of TSS validated by both our data and PRO-cap data versus TSS validated only by PRO-cap data, we sorted both kinds of TSS (PRO-cap 96 hpi and dRNA-seq 96 hpi) in descending order. Then, from top to bottom, we picked TSS pairs of both sets with similar read counts (+/-5% difference per pair). TSS, which were in close proximity (1,000 bp) to an already picked TSS were discarded (i.e. if multiple TSS were in close proximity, only the one with the highest read count was considered). Translation rates of TSS were calculated by using the codon-counts calculated using the tool PRICE ^5^ in the Ribo-seq data treated with either harringtonine (HRT) or lactimidomycin (LTM), respectively. Subsequently, for each TSS, the translational efficiency was calculated by dividing the codon count of the codon with the highest codon count within a 1kb downstream window by the read count of the respective TSS.

## Data availability

The dRNA-seq and STRIPE-seq data produced in this study are available at GEO (accession number GSE191299). Already published data used in this study can be found at the following sites: PRO(cap)-seq (GEO; accession number GSE113394), PacBio and MinION (European Nucleotide Archive; accession number PRJEB25680).

## Code availability

The gedi toolkit, which was used for mapping and most of the analysis steps, is available on GitHub (https://github.com/erhard-lab/gedi). iTiSS, which is a module for gedi is available separately on GitHub (https://github.com/erhard-lab/iTiSS). The tool MetagenePlot, which is also a module for gedi, can be found on GitHub as well (https://github.com/erhard-lab/MetagenePlot). The source code of the gedi toolkit itself is freely available on GitHub (https://github.com/erhard-lab/gedi). The source code of all additional custom scripts generated for generating the Figures, tables and analyzing the data in general can be found at Zenodo (https://doi.org/10.5281/zenodo.5801030).

## Extended Data Figures

**Extended Data Fig. 1**

**A**. Genome browser screenshot for the UL112-UL122 region of the HCMV genome. The upper half shows the plus strand, the lower half the minus strand. Canonical ORFs are depicted in green. Data shown is the total 5’ read count over all time points for dRNA-seq (replicate 2) and STRIPE-seq (replicate 1), respectively.

**B**,**C**. Pairwise correlation analysis of the read counts per nucleotide in the HCMV genome for individual replicates per time point in the dRNA-seq (**B**) or STRIPE-seq (**C**) data. Pearson correlation coefficients (r) are indicated.

**D**. TSS enrichment (read count at the TSS +/-1 bp divided by the read count 100 bp downstream of the TSS) at annotated cellular TSS in the dRNA-seq (red) and STRIPE-seq (blue) sample.

**E**,**F**. Comparison of long-lived (red) and short-lived (blue) RNAs based on the predicted new-to-total RNA ratios (NTRs) in the dRNA-seq (**E**) and STRIPE-seq (**F**) data. NTRs were estimated using GRAND-SLAM ^17^. The NTRs for GAPDH and MYC are indicated.

**G**,**H**. The ratios of all occurring nucleotide mismatches in our dRNA-seq (**G**) and STRIPE-seq (**H**) experiment, respectively. Mismatch ratios were calculated using GRAND-SLAM ^17^.

**Extended Data Fig. 2**

**A**,**B**. Sequence logos of the 100 most strongly expressed TSS of each time point based on newly synthesized RNA (A) or total RNA (B), respectively. For cellular genes, only the mock (0 hpi) data is shown. Sequence logos range from 100 bp upstream to 100 bp downstream of the respective TSS.

**C**. Log_2_ odds of observed vs. expected occurrences of correctly positioned promoter elements (see Extended Data Tab. 3). The critical region (p<0.01, binomial test) is indicated as a dashed line.

**D**. Results of the differential motif analysis using MEME. Shown are the three top scoring (lowest E-value) motifs for the TATA- and TATT-associated TSS, respectively.

**E**. The cumulative frequency of the indicated di-nucleotides directly downstream of all TATA and TATT-boxes in the HCMV genome, respectively, based on the maximum read count in their respective expected Inr-region.

**F**. All significantly enriched motifs found using MEME in the HCMV or cellular genome, respectively.

**Extended Data Fig. 3**

**A**,**B**. Abundance profiles for all TATA-TSS with at least 20 reads, TLS>150% (**A**) or TLS<30% (**B**) and without a canonical ORF located within 1 kb downstream for total RNA (solid line) and newly synthesized RNA (dashed line). GCV treated samples are indicated as a triangle. For each TSS, the upper triangle depicts total RNA the lower newly synthesized RNA.

**C**. (Top) Locations of viral TATA- and TATT-boxes relative to the TSS (identical to Fig. 2C without the cellular data). (Bottom) TLS scores and locations of TATA-associated TSS with ≥20 reads.

**D**. Boxplots showing the alternative “‘True Late Score” (ratio of normalized expression values 24hpi vs. 72hpi) for all TATA- or TATT-TSS, TSS with ≥10 reads, TSS with ≥ 20 reads and TSS with ≥ 50 reads, respectively.

**Extended Data Fig. 4**

**A**. Translational efficiency (translation / transcription) for TATA- and TATT-associated TSS identified at 2 hpi. Only TSS upstream of canonical large ORFs were. Translation strength was estimated from the Ribo-seq codon count at the start codon of the respective ORF in the LTM treated sample. Statistical significance according to a Mann-Whitney-U test is indicated (**: p<1.0^-2^; the p-values were: 6 hpi, p = 0.001; 72 hpi, p = 0.24).

**B**. Change in translational efficiency between 6 hpi and 72 hpi for TATA and TATT associated TSS identified at 2 hpi. Only TSS upstream of canonical ORFs were selected. Translation strength was estimated as in A. Statistical significance according to a Mann-Whitney-U test is indicated (**: p<1.0^-2^; p-value for 72/6 hpi: 0.006).

**C**. Scatterplot comparing the fold changes between 48 and 12 hpi (x-axis) and 96 and 48 hpi (y-axis) for protein abundances. Proteins are colored based on their assigned Tp-cluster. 2 outlier points with x or y values >7 from Tp 4&5 are omitted.

**D**. The mean abundance score for each temporal RNA (Tr) profile. Reads were clustered using the KMeans++ algorithm on the 6, 12, 48, 72 and 96 hour timepoint with k=5 using only the dominant TSS per ORF (n=162). Subsequently, the remaining TSS (n=354) were assigned to their most similar cluster.

**Extended Data Fig. 5**

**A**. Sequence logo for the top 100 most strongly transcribed (top) and all TSS (bottom) validated by our datasets (i.e. found in dRNA-seq and STRIPE-seq), but not by PRO-cap.

**B**. Heatmaps showing the total 5’-read counts over all time points in the dRNA-seq data (left) and the total 5’-read counts at 96 hpi in the PRO-cap data (right) around all TATT-motifs found in the HCMV genome. The TATT-motifs as well as the expected region of the TSS are indicated. The heatmap is sorted according to the number of reads inside the Inr region of the dRNA-seq data. The threshold of ≥10 reads within the Inr region in the dRNA-seq dataset is indicated.

**C**. Comparison of the translation rates downstream of TSS identified in both data sets (PRO-cap and dRNA-seq) or only in dRNA-seq inferred from the harringtonine (HRT) or lactimidomicin (LTM) treated Ribo-seq samples. dRNA-seq only TSS were selected to match the expression strength of TSS identified in both data sets. The differences were not significant (Mann-Whitney-U test; p-values: HRT (Rep 1)=0.89, HRT (Rep 2)=0.74, LTM (Rep 1)=0.37, LTM (Rep 2)=0.39).

**D**. Comparison of the translation rates downstream of TSS identified in both data sets (PRO-cap and STRIPE-seq) or only in STRIPE-seq inferred from the harringtonine (HRT) or lactimidomicin (LTM) treated Ribo-seq samples. STRIPE-seq only TSS were selected to match the expression strength of TSS identified in both data sets. The differences are not significant (Mann-Whitney-U test; p-values: HRT (Rep 1)=0.97, HRT (Rep 2)=0.77, LTM (Rep 1)=0.47, LTM (Rep 2)=0.47).

**E**. UL83/UL84 locus in the genome browser. TSS giving rise to transient and stable RNAs are indicated. Tracks show from top to bottom: The dRNA-seq 5’-counts at 96 hpi on the plus strand, the 5’-counts at 96 hpi of the PROcap-seq data on the plus strand, the dRNA-seq 5’-counts at 96 hpi on the minus strand and finally the 5’-counts at 96 hpi of the PROcap-seq data on the minus strand.

## Extended Data Tables

**Extended Data Tab. 1**

Conversion rates and error rates determined by GRAND-SLAM

**Extended Data Tab. 2**

The top 500 TSS most strongly expressed TSS found per timepoint. For each timepoint, TSS are categorized into 5 bins, with each holding 100 TSS. Bin 1 contains the 100 most strongly transcribed TSS bin 2 the second 100 most strongly transcribed, and so forth.

**Extended Data Tab. 3**

For each TSS in either human or HCMV the distance of the respective promoter sequence to the TSS is indicated. Negative implies upstream and positive downstream. The TSS has position 0.

**Extended Data Tab. 4**

For each timepoint all transcription factors identified by TFM-Explorer ^29^ in the top 500 TSS are shown.

## References

1. Yinon, Y. et al. Cytomegalovirus Infection in Pregnancy. J. Obstet. Gynaecol. Canada 32, 348–354 (2010).

2. Stagno, S. et al. Primary Cytomegalovirus Infection in Pregnancy: Incidence, Transmission to Fetus, and Clinical Outcome. JAMA 256, 1904–1908 (1986).

3. Pultoo, A., Jankee, H., Meetoo, G., Pyndiah, M. N. & Khittoo, G. Detection of cytomegalovirus in urine of hearing-impaired and mentally retarded children by PCR and cell culture. J. Commun. Dis. 32, 101–108 (2000).

4. Stern-Ginossar, N. et al. Decoding Human Cytomegalovirus. Science 338, 1088–1093 (2012).

5. Erhard, F. et al. Improved Ribo-seq enables identification of cryptic translation events. Nat. Methods 15, 363–366 (2018).

6. Hiroki, I. et al. The Human Cytomegalovirus Gene Products Essential for Late Viral Gene Expression Assemble into Prereplication Complexes before Viral DNA Replication. J. Virol. 85, 6629–6644 (2011).

7. Li, M. et al. Cytomegalovirus late transcription factor target sequence diversity orchestrates viral early to late transcription. PLOS Pathog. 17, e1009796 (2021).

8. Gruffat, H., Marchione, R. & Manet, E. Herpesvirus Late Gene Expression: A Viral-Specific Pre-initiation Complex Is Key. Front. Microbiol. 7, 869 (2016).

9. Sarisky, R. T. & Hayward, G. S. Evidence that the UL84 gene product of human cytomegalovirus is essential for promoting oriLyt-dependent DNA replication and formation of replication compartments in cotransfection assays. J. Virol. 70, 7398– 7413 (1996).

10. Malone, C. L., Vesole, D. H. & Stinski, M. F. Transactivation of a human cytomegalovirus early promoter by gene products from the immediate-early gene IE2 and augmentation by IE1: mutational analysis of the viral proteins. J. Virol. 64, 1498– 1506 (1990).

11. Weekes, M. P. et al. Quantitative temporal viromics: an approach to investigate host-pathogen interaction. Cell 157, 1460–1472 (2014).

12. Parida, M. et al. Nucleotide Resolution Comparison of Transcription of Human Cytomegalovirus and Host Genomes Reveals Universal Use of RNA Polymerase II Elongation Control Driven by Dissimilar Core Promoter Elements. MBio 10, (2019).

13. Li, M. et al. Human cytomegalovirus IE2 drives transcription initiation from a select subset of late infection viral promoters by host RNA polymerase II. PLOS Pathog. 16, e1008402 (2020).

14. Policastro, R. A., Raborn, R. T., Brendel, V. P. & Zentner, G. E. Simple and efficient profiling of transcription initiation and transcript levels with STRIPE-seq. Genome Res. (2020). doi:10.1101/gr.261545.120

15. Sharma, C. M. & Vogel, J. Differential RNA-seq: the approach behind and the biological insight gained. Curr. Opin. Microbiol. 19, 97–105 (2014).

16. Whisnant, A. W. et al. Integrative functional genomics decodes herpes simplex virus 1. Nat. Commun. 11, 2038 (2020).

17. Jürges, C., Dölken, L. & Erhard, F. Dissecting newly transcribed and old RNA using GRAND-SLAM. Bioinformatics 34, i218.-i226 (2018).

18. Erhard, F. et al. scSLAM-seq reveals core features of transcription dynamics in single cells. Nature 571, 419–423 (2019).

19. Jürges, C. S., Dölken, L. & Erhard, F. Integrative transcription start site identification with iTiSS. Bioinformatics (2021). doi:10.1093/bioinformatics/btab170

20. Balázs, Z., Tombácz, D., Szűcs, A., Snyder, M. & Boldogkői, Z. Dual Platform Long-Read RNA-Sequencing Dataset of the Human Cytomegalovirus Lytic Transcriptome Frontiers in Genetics 9, 432 (2018).

21. Perera, L. P. The TATA Motif Specifies the Differential Activation of Minimal Promoters by Varicella Zoster Virus Immediate-early Regulatory Protein IE62*. J. Biol. Chem. 275, 487–496 (2000).

22. Sambucetti, L. C., Cherrington, J. M., Wilkinson, G. W. & Mocarski, E. S. NF-kappa B activation of the cytomegalovirus enhancer is mediated by a viral transactivator and by T cell stimulation. EMBO J. 8, 4251–4258 (1989).

23. Cherrington, J. M. & Mocarski, E. S. Human cytomegalovirus ie1 transactivates the alpha promoter-enhancer via an 18-base-pair repeat element. J. Virol. 63, 1435–1440 (1989).

24. Haberle, V. & Stark, A. Eukaryotic core promoters and the functional basis of transcription initiation. Nat. Rev. Mol. Cell Biol. 19, 621–637 (2018).

25. Vo Ngoc, L. et al. The punctilious RNA polymerase II core promoter. 31, 1289–1301 (Cold Spring Harbor Laboratory Press, 2017).

26. Bailey, T. L. & Elkan, C. Fitting a mixture model by expectation maximization to discover motifs in biopolymers. Proceedings. Int. Conf. Intell. Syst. Mol. Biol. 2, 28–36 (1994).

27. Blake, M. C., Jambou, R. C., Swick, A. G., Kahn, J. W. & Azizkhan, J. C. Transcriptional initiation is controlled by upstream GC-box interactions in a TATAA-less promoter. Mol. Cell. Biol. 10, 6632–6641 (1990).

28. Parry, T. J. et al. The TCT motif, a key component of an RNA polymerase II transcription system for the translational machinery. Genes Dev. 24, 2013–2018 (2010).

29. Tonon, L., Touzet, H. & Varré, J.-S. TFM-Explorer: mining cis-regulatory regions in genomes. Nucleic Acids Res. 38, W286–W292 (2010).

30. McFarlane, S., Nicholl, M. J., Sutherland, J. S. & Preston, C. M. Interaction of the human cytomegalovirus particle with the host cell induces hypoxia-inducible factor 1 alpha. Virology 414, 83–90 (2011).

31. Wise, L. M., Xi, Y. & Purdy, J. G. Hypoxia-Inducible Factor 1α (HIF1α) Suppresses Virus Replication in Human Cytomegalovirus Infection by Limiting Kynurenine Synthesis. MBio 12, (2021).

32. Hagemeier, C., Walker, S., Caswell, R., Kouzarides, T. & Sinclair, J. The human cytomegalovirus 80-kilodalton but not the 72-kilodalton immediate-early protein transactivates heterologous promoters in a TATA box-dependent mechanism and interacts directly with TFIID. J. Virol. 66, 4452–4456 (1992).

33. Marcinowski, L. et al. Real-time Transcriptional Profiling of Cellular and Viral Gene Expression during Lytic Cytomegalovirus Infection. PLoS Pathog 8, e1002908 (2012).

34. Amati, B., Frank, S. R., Donjerkovic, D. & Taubert, S. Function of the c-Myc oncoprotein in chromatin remodeling and transcription. Biochim. Biophys. Acta Rev. Cancer 1471, M135–M145 (2001).

35. Davis, Z. H. et al. Global Mapping of Herpesvirus-Host Protein Complexes Reveals a Transcription Strategy for Late Genes. Mol. Cell 57, 349–360 (2015).

36. Terhune, S. S., Schroer, J. & Shenk, T. RNAs Are Packaged into Human Cytomegalovirus Virions in Proportion to Their Intracellular Concentration. J. Virol. 78, 10390–10398 (2004).

37. Bresnahan, W. A. & Shenk, T. A subset of viral transcripts packaged within human cytomegalovirus particles. Science (80-.). 288, 2373–2376 (2000).

38. Preker, P. et al. RNA exosome depletion reveals transcription upstream of active human promoters. Science 322, 1851–1854 (2008).

39. Herzog, V. A. et al. Thiol-linked alkylation of RNA to assess expression dynamics. Nat. Methods 14, 1198–1204 (2017).

40. Hinnebusch, A. G. & Lorsch, J. R. The mechanism of eukaryotic translation initiation: new insights and challenges. Cold Spring Harb. Perspect. Biol. 4, a011544 (2012).

41. Sinclair, J. Chromatin structure regulates human cytomegalovirus gene expression during latency, reactivation and lytic infection. Biochim. Biophys. Acta 1799, 286–295 (2010).

42. Rahl, P. B. & Young, R. A. MYC and transcription elongation. Cold Spring Harb. Perspect. Med. 4, (2014).

43. Saldi, T., Cortazar, M. A., Sheridan, R. M. & Bentley, D. L. Coupling of RNA Polymerase II Transcription Elongation with Pre-mRNA Splicing. Journal of Molecular Biology 428, 2623–2635 (2016).

44. Hein, M. Y. & Weissman, J. S. Functional single-cell genomics of human cytomegalovirus infection. Nat. Biotechnol. 1–11 (2021). doi:10.1038/s41587-021-01059-3

45. Caragliano, E. et al. Human Cytomegalovirus Forms Phase-Separated Compartments at Viral Genomes to Facilitate Viral Replication. SSRN Electron. J. (2021). doi:10.2139/ssrn.3871395

46. Winkler, M., Rice, S. A. & Stamminger, T. UL69 of human cytomegalovirus, an open reading frame with homology to ICP27 of herpes simplex virus, encodes a transactivator of gene expression. J. Virol. 68, 3943–3954 (1994).

47. Wang, X. et al. Herpes simplex virus blocks host transcription termination via the bimodal activities of ICP27. Nat. Commun. 11, 293 (2020).

48. Ruiz, J. C., Hunter, O.V & Conrad, N. K. Kaposi’s sarcoma-associated herpesvirus ORF57 protein protects viral transcripts from specific nuclear RNA decay pathways by preventing hMTR4 recruitment. PLOS Pathog. 15, e1007596 (2019).

49. Le-Trilling, V. T. K. et al. The Human Cytomegalovirus pUL145 Isoforms Act as Viral DDB1-Cullin-Associated Factors to Instruct Host Protein Degradation to Impede Innate Immunity. Cell Rep. 30, 2248-2260.e5 (2020).

50. Bolger, A. M., Lohse, M. & Usadel, B. Trimmomatic: a flexible trimmer for Illumina sequence data. Bioinformatics 30, 2114–2120 (2014).

51. Langmead, B. & Salzberg, S. L. Fast gapped-read alignment with Bowtie 2. Nat. Methods 9, 357–359 (2012).

52. Dobin, A. et al. STAR: ultrafast universal RNA-seq aligner. Bioinformatics 29, 15–21 (2013).

53. Li, H. et al. The Sequence Alignment/Map format and SAMtools. Bioinformatics 25, 2078–2079 (2009).

54. Smith, T. S., Heger, A. & Sudbery, I. UMI-tools: Modelling sequencing errors in Unique Molecular Identifiers to improve quantification accuracy. Genome Res. (2017). doi:10.1101/gr.209601.116

55. Wu, T. D. & Watanabe, C. K. GMAP: a genomic mapping and alignment program for mRNA and EST sequences. Bioinformatics 21, 1859–1875 (2005).

56. Davis, M. P. A., van Dongen, S., Abreu-Goodger, C., Bartonicek, N. & Enright, A. J. Kraken: a set of tools for quality control and analysis of high-throughput sequence data. Methods 63, 41–49 (2013).

57. Langmead, B., Trapnell, C., Pop, M. & Salzberg, S. L. Ultrafast and memory-efficient alignment of short DNA sequences to the human genome. Genome Biol. 10, R25 (2009).

58. Wagih, O. ggseqlogo: A ‘ggplot2’ Extension for Drawing Publication-Ready Sequence Logos. (2017).

59. Arthur, D. & Vassilvitskii, S. K-Means++: The Advantages of Careful Seeding. in Proceedings of the Eighteenth Annual ACM-SIAM Symposium on Discrete Algorithms 1027–1035 (Society for Industrial and Applied Mathematics, 2007).

