## Supplementary figures and images for "Multi-omics reveals principles of gene regulation and pervasive non-productive transcription in the human cytomegalovirus genome"

### Supplementary Figure 1

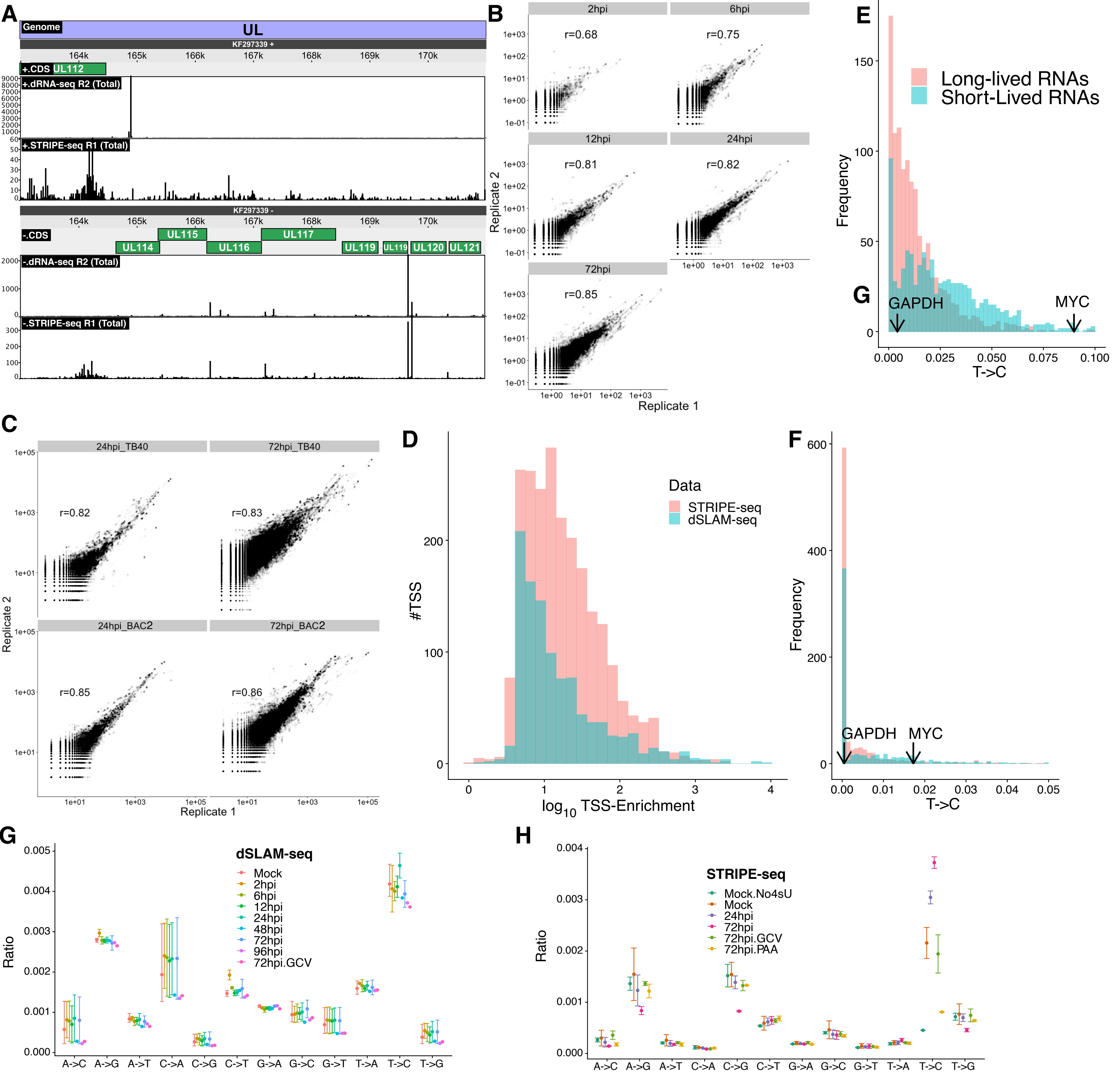

### Supplementary Figure 2

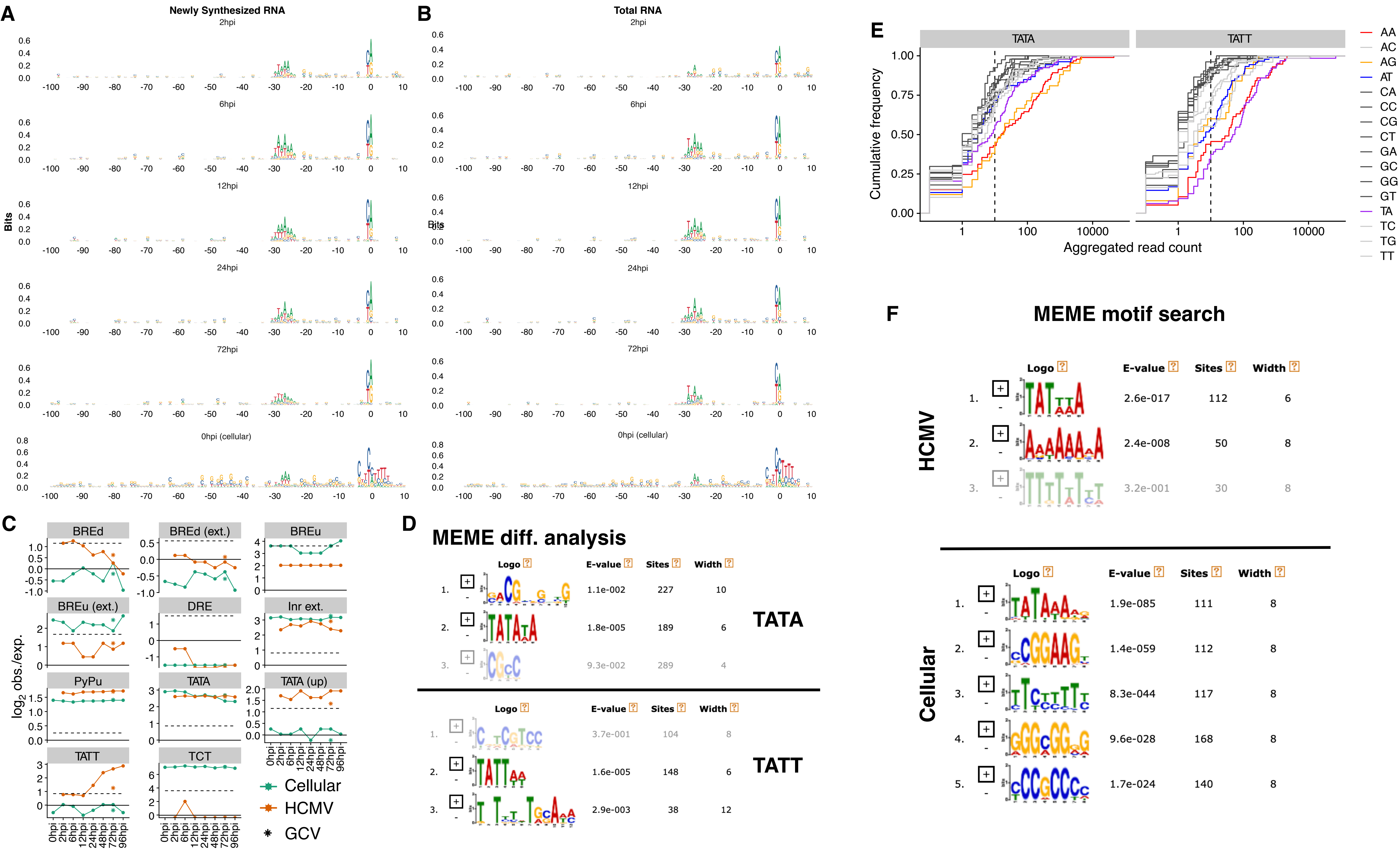

### Supplementary Figure 3

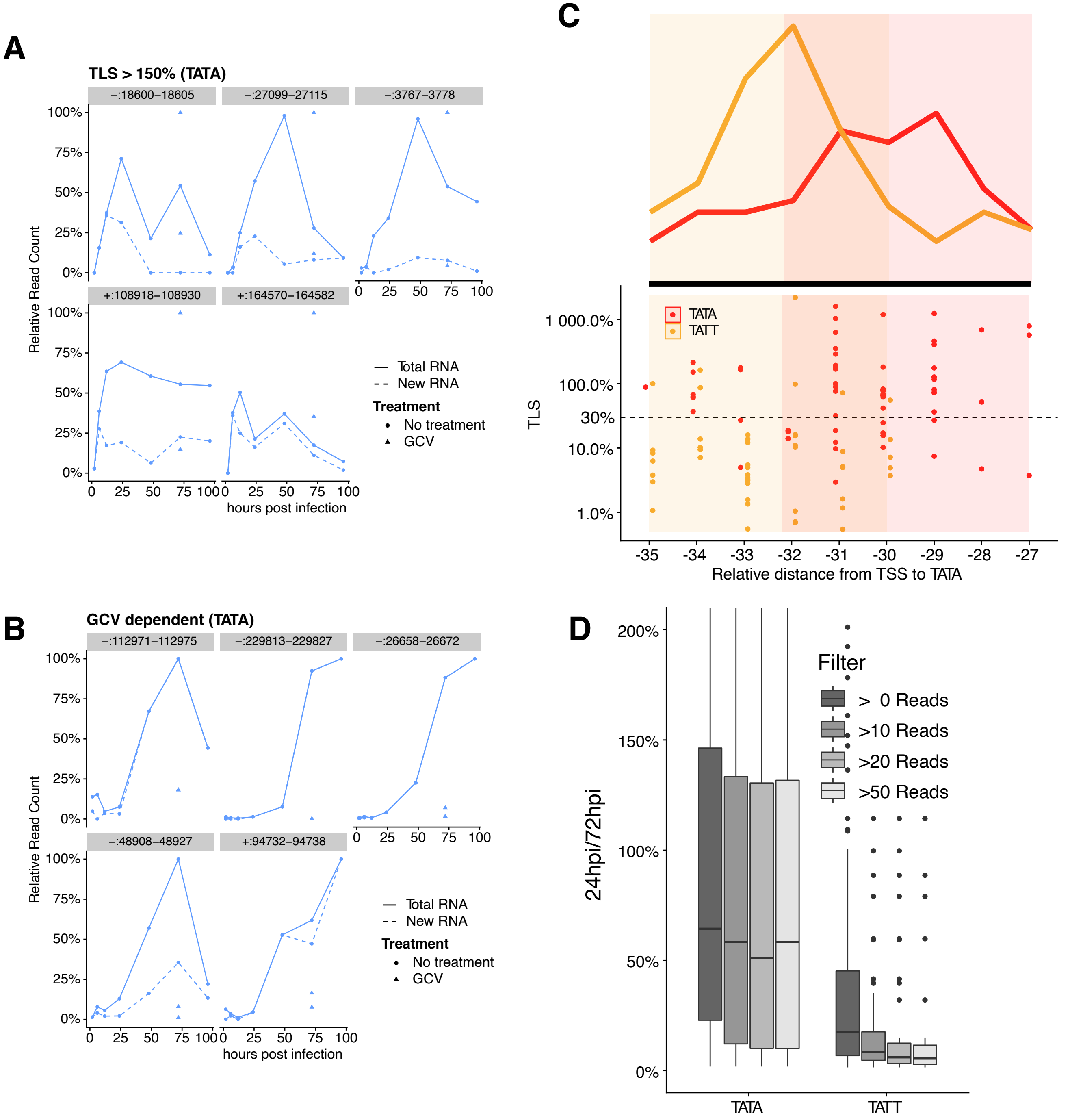

### Supplementary Figure 4

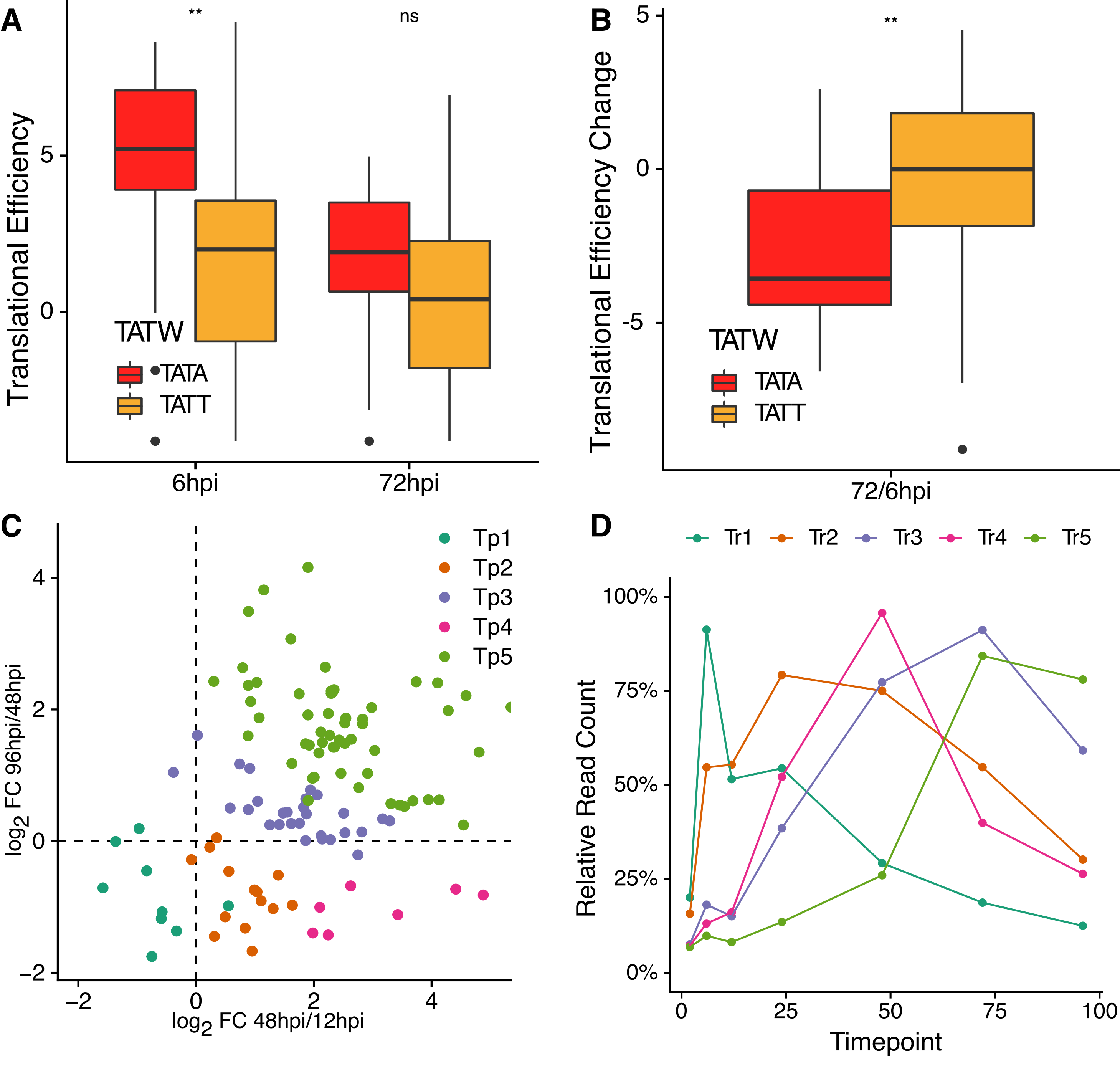

### Supplementary Figure 5

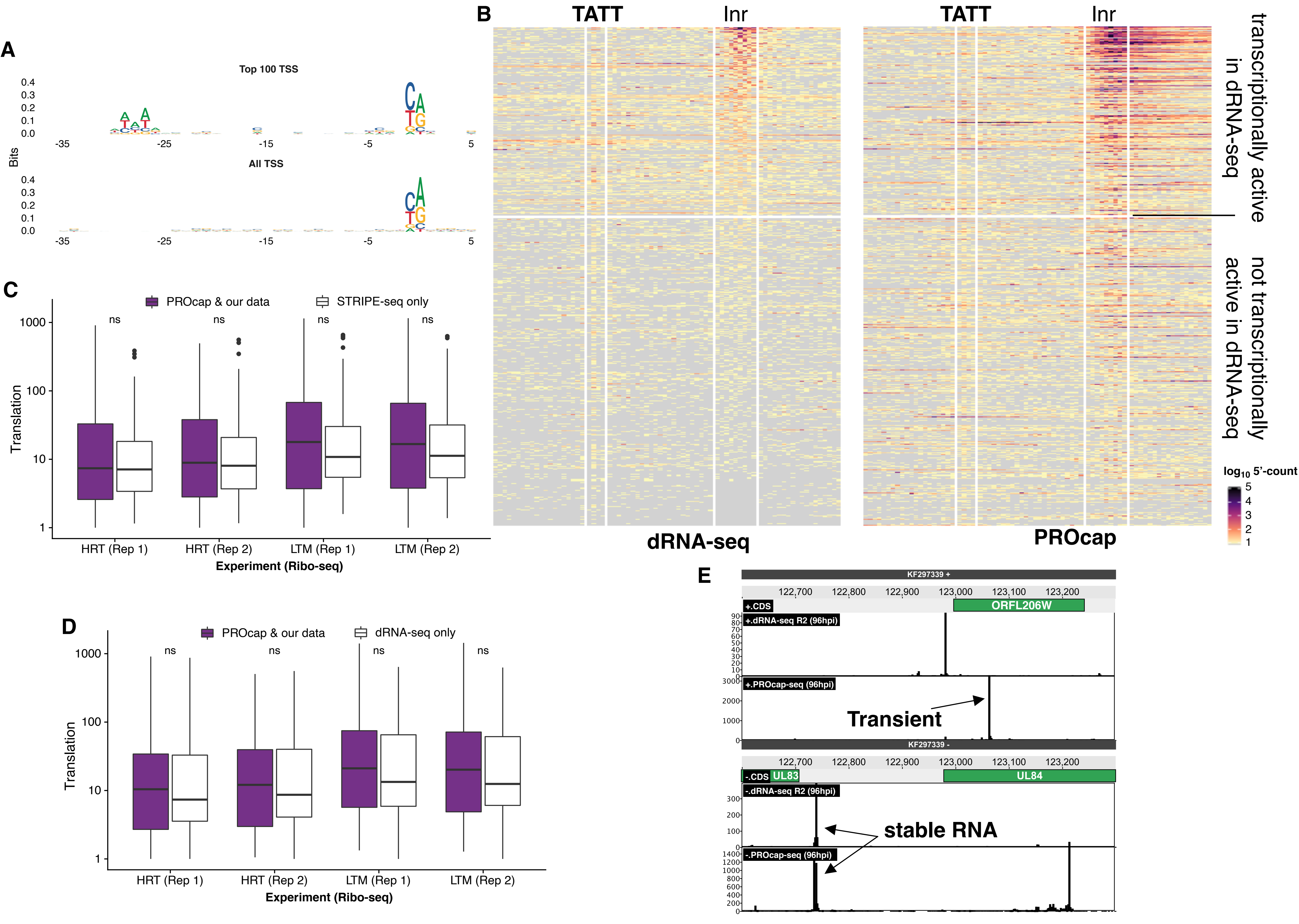
